# Molecular Architecture of the Human GCN1–ABCF3 Ribosome Collision Sensor

**DOI:** 10.64898/2026.09.19.752920

**Authors:** Viswanathan Chandrasekaran, Ritu Gupta, Mary T. Angani, Talal F. M. Haddad, Christopher G. Tate, V. Ramakrishnan, Alan G. Hinnebusch

## Abstract

The Integrated Stress Response is a critical eukaryotic signaling pathway that maintains cellular proteostasis. During amino acid scarcity, the yeast kinase Gcn2 (GCN2 in humans) and its activators Gcn1/Gcn20 phosphorylate eIF2α to reprogram translation, but how GCN2 senses nutrient stress in mammals is not known. Ribosome collisions serve as a major physiological trigger. Here we report a cryo-EM structure of human GCN1 bound to collided di-ribosomes. GCN1 specifically recognizes and rigidifies the flexible interface of the collided di-ribosome. Binding is mediated by contacts with both ribosomal P-stalks and a conserved segment that “pinches” the beak of the 40S subunit of the trailing ribosome. Functional assays confirm that multivalent Gcn1 contacts with collided ribosomes are essential for full Gcn2 activation in stressed yeast cells. Using nanobody-based proteomics in cells deprived of prolyl-tRNA, we identify ABCF3 as the primary mammalian ortholog of yeast Gcn20 recruited to ribosomes during amino acid starvation.

## Main Text

The Integrated Stress Response (ISR) is a universal eukaryotic signaling network that enables cellular adaptation to environmental and intrinsic stressors^1, 2, 3, 4^. In mammals, the ISR is orchestrated by four distinct specialized kinases, each signaling for a specific type of physiological stress. GCN2 (General Control Non-derepressible 2) senses amino acid starvation. PKR (Protein Kinase R) detects cytosolic double-stranded RNA, a hallmark of viral infection. PERK (PKR-like ER Kinase) monitors protein folding stress in the endoplasmic reticulum and HRI (Heme-Regulated Inhibitor) responds to heme deficiency and oxidative stress. Despite their diverse sensory domains, these four kinases converge on a single, highly conserved regulatory step: phosphorylation of the alpha subunit of eukaryotic initiation factor 2 (eIF2α) at serine 51.

The eIF2 complex is a heterotrimeric GTPase that plays a critical role in delivering initiator methioninyl-tRNA (Met-tRNA^i,Met^) to the 40S ribosomal subunit to form a 43S pre-initiation complex (reviewed in ^5^). This delivery is dependent on the nucleotide state of eIF2; it must be bound to GTP to possess high affinity for the initiator tRNA. Following successful identification of the start codon and GTP hydrolysis, eIF2 is released in a “spent” GDP-bound state. Because eIF2 has a high affinity for GDP, it cannot spontaneously exchange GDP for GTP, instead relying on eIF2B, a large decameric complex that serves as the essential Guanine Nucleotide Exchange Factor (GEF) for eIF2^6^. Under normal conditions, eIF2B ensures a steady supply of active eIF2-GTP to maintain high rates of protein synthesis. However, once phosphorylated by one of the four ISR kinases, phosphorylated eIF2 (p-eIF2) transforms from substrate to potent competitive inhibitor of eIF2B. Because the cellular concentration of eIF2 significantly exceeds that of eIF2B, even a modest increase in p-eIF2 levels leads to total sequestration of eIF2B in an inactive complex^6^. This elicits a global downregulation of cap-dependent translation. Paradoxically, this global suppression is accompanied by enhanced translation of specific stress-adaptive mRNAs, such as encoding Gcn4 in yeast and ATF4 in mammals, which contain upstream open reading frames (uORFs)^7, 8, 9^. These factors then initiate a transcriptional program aimed at restoring cellular balance. If the damage is terminal, however, the ribotoxic stress response is engaged and leads ultimately to apoptosis^10^.

Following standard nomenclature guidelines, human proteins are hereafter denoted in all uppercase letters (GCN1, GCN2) and yeast proteins are denoted with an initial capital letter (Gcn1, Gcn2). Where generic functions applicable to both organisms are discussed, GCN1/GCN2/the GCN2 family is used.

Among the ISR kinases, the GCN2 family represents the most evolutionarily conserved branch, with orthologs spanning from budding yeast to humans^7, 11^. This conservation likely underscores the fundamental importance of sensing amino acid availability. From yeast to human, GCN2 has a conserved architecture^12, 13^ (Figure S1A) comprising an N-terminal RWD domain that is essential for mediating protein-protein interaction with its activator GCN1, a regulatory pseudokinase domain that modulates the activity of the neighbouring latent kinase domain, which is the catalytic domain responsible for eIF2α phosphorylation and GCN2 autophosphorylation, a histidyl-tRNA synthetase-related (HisRS-like) domain that is historically thought to be the primary sensor of uncharged tRNAs and a C-terminal dimerization domain that is critical for forming the active dimeric state of GCN2^13, 14, 15, 16, 17^. The prevailing model suggested that the HisRS-like domain is directly bound to uncharged tRNAs that accumulate during amino acid starvation, triggering a conformational change that activated the kinase. There was strong evidence however that yeast Gcn2 is activated while tethered to translating ribosomes in association with Gcn1, presumably by uncharged tRNA paired with a “hungry codon” for the cognate limiting acid positioned in the ribosomal A site^14, 15^, akin to the activation of bacterial RelA in the stringent response^18^. Structural data indicates that yeast Gcn1 binds to disomes formed by collision between a stalled ribosome and a trailing elongating ribosome^19^. Thus, GCN2 may be activated in amino acid-starved cells on collided ribosomes produced by stalling of the leading ribosome with uncharged tRNA cognate to the hungry codon in the A site.

GCN2 can also be activated by stalled ribosomes in the absence of uncharged tRNA. Such activation has been seen in mutant mammalian cells with reduced cognate tRNA for a particular amino acid and lacking a ribosome rescue factor^20^, in the presence of translation elongation inhibitors or agents that damage mRNA in yeast, *Neurospora*, and mammalian cells^10, 21, 22, 23^. In untreated *Neurospora* cells, elongating ribosomes paused at rare codons can also activate Gcn2^23^. *In vitro*, mammalian GCN2 can be activated by ribosomes or by the isolated P stalk of the 60S subunit. This activation occurs independently of tRNA but requires the HisRS-like domain of GCN2. The ribosomal P stalk, which comprises the proteins uL10 and two P1/P2 heterodimers, interacts with the elongation GTPases eEF1A and eEF2. It has been suggested that the P stalk becomes accessible to GCN2 when the A site lacks an incoming aminoacylated tRNA bound to elongation factor eEF1A^13^. Consistently, P stalk mutations impair GCN2 activation in amino acid-starved human cells^24^. Interestingly, the P stalk P1/P2 heterodimer was shown to be essential for activation of yeast Gcn2 in cells harboring ribosomes stalled with an empty A site, but not in amino acid-starved cells where uncharged tRNA might substitute for P1/P2 proteins as activating ligand^25^. Recent biochemical studies in mammalian cell extracts confirmed that uncharged tRNAs are neither necessary nor sufficient for GCN2 activation, and that GCN1-dependent activation requires formation of a GCN1-bound collided ribosome queue capable of recruiting GCN2 as well as ribosome-interaction moieties within GCN2. Addition of uncharged tRNA cognate to the hungry codon of the stalled ribosomes strongly enhanced GCN2 activation in this system^26^, consistent with distinct activation mechanisms conferring different levels of GCN2 activation.

Thus, the following model emerges for GCN2 activation in mammalian or yeast cells deprived of a specific amino acid: When the concentration of the corresponding aminoacyl-tRNA drops, ribosomes stall at the hungry codon awaiting the cognate aminoacylated tRNA. As translation initiation continues upstream, subsequent ribosomes continue to elongate until they collide with the stalled leading ribosome. These collided di-ribosomes (disomes) create a unique structural “molecular signature” that is distinct from the architecture of an actively translating polysome^27, 28^. Indeed, such collided ribosomes serve as a signalling platform in several other important cellular contexts. In mammals, sensors such as ZNF598 and ZAK recognize these collisions to trigger ribosome-associated quality control (RQC) or ribotoxic stress signaling (RSR)^27, 29^. In the case of GCN2, activation requires its highly conserved binding partner and activator, GCN1^15^. GCN1 is a large, 300 kDa protein comprising HEAT repeats (Figure S1A) and yeast Gcn1 exists as a constitutive heterodimer with the ATPase Gcn20, whose activating function is ill-defined^30^. Binding of GCN1 to the collided disome enables recruitment of GCN2 in a manner that positions the HisRS-like domain near the empty A site of the stalled ribosome, allowing allosteric activation of the GCN2 kinase domain by P-stalk proteins or its hyperactivation by cognate uncharged tRNA bound in the A site.

While a cryo-EM structure of yeast Gcn1 bound to a disome^19^ provided an initial framework for Gcn2 activation on collided stalled ribosomes, the structural biology of the mammalian response to amino acid starvation remains unresolved. It is currently unclear how GCN1, GCN2 and the mammalian homolog of Gcn20 recognize the specific architecture of the mammalian collided disome or how these interactions are coupled to kinase activation.

ABCF proteins constitute a distinct sub-family of the ATP-binding cassette (ABC) superfamily. Unlike canonical ABC transporters, members of the ABCF family lack transmembrane domains and operate instead as soluble cytoplasmic factors primarily dedicated to the regulation of protein synthesis and translational stress responses^31^. Structurally, these proteins feature two highly conserved nucleotide-binding domains (NBDs) connected by an essential regulatory linker region, alongside specialized terminal extensions that facilitate direct, high-affinity interactions with the E site of the ribosome^31^. In yeast, the ABCF protein Gcn20 forms a stable complex with Gcn1 to mediate the downstream activation of Gcn2 during amino acid deprivation^19, 32^. Humans possess three distinct paralogs—ABCF1, ABCF2, and ABCF3—among which ABCF3 shares the highest sequence homology and architectural similarity with yeast Gcn20^33^. While mammalian ABCF proteins have been generally implicated in translation initiation, mRNA translation control, and ribosome quality control, their precise biochemical recruitment, structural configuration, and functional integration into the human GCN1-dependent ribosome collision-sensing machinery remain poorly understood.

In this study, we set out to investigate the molecular principles of ribosome recognition by GCN proteins during amino acid starvation. We asked whether ribosome collisions are necessary for GCN1 recruitment and GCN2 activation. We aimed to understand the structural basis of how mammalian GCN1 detects ribosome collisions to trigger phosphorylation of eIF2α by GCN2. Structural insights gained from studying mammalian GCN proteins could be readily tested *in vivo* using mutagenesis studies in the conserved yeast system. We also sought to understand the differences, if any, between the yeast and mammalian systems, including the identities of factors that associate with GCN1 and GCN2 and whether the mechanism of GCN2 activation is conserved.

We present a cryo-EM structure of human GCN1 in complex with the collided disome. Our data reveal a “clamp” mechanism by which GCN1 rigidifies the flexible disome interface and interacts with the ribosomal P-stalk. Using yeast *in vivo* assays, we show that Gcn1 recognition of ribosome collisions is necessary for full Gcn2 activation. Through nanobody-based proteomics in human cells, we identify ABCF3 as the mammalian ortholog of yeast Gcn20 and show that it associates with GCN1 during amino acid starvation. Together, these findings provide a structural and mechanistic blueprint for the mammalian amino acid stress response, rationalizing decades of genetic data and defining the molecular and structural determinants of GCN2 activation.

## Results

### Reconstitution and structure determination of Human GCN1–Disome Complexes

To assemble ribosome-GCN protein complexes, we first reconstituted ribosome stalling and collisions in a rabbit reticulocyte lysate (RRL)-based in vitro translation system^34^. To stall ribosomes in a way that facilitates ribosome collisions, we employed a trick that we used previously to stall polyribosomes on globin mRNAs (Figure S1B)^27^. A dominant-negative eukaryotic release factor 1 (eRF1^AAQ^) was used to arrest only the leading ribosomes that have reached stop codons on alpha-, or beta-globin transcripts^35^. Upon binding terminating ribosomes that have reached stop codons, wild type eRF1 utilizes a conserved GGQ motif in its M domain to hydrolyze the nascent chain^36^. Mutant eRF1^AAQ^ also accommodates into the A site, but fails to hydrolyze the completed globin polypeptide, instead remaining in the A site and preventing termination, ribosome splitting and recycling^35^. Trailing ribosomes on these polysomes, however, continue to elongate unencumbered until they collide with the stalled ribosome or join the queue of collided ribosomes.

We expressed and purified human GCN2 and GCN1 recombinantly with N-terminal Twin-Strep-tags using an insect cell-baculovirus system^13^ (Figure S1C). We then performed in vitro translation in the presence of pure recombinant eRF1^AAQ^, Twin-Strep-tagged human GCN2 and GCN1 and human EDF1, which is known to coordinate signaling responses to ribosome collisions in a variety of contexts, including the ISR, by binding at the interface between the stalled and collided ribosomes^19, 37, 38^. Following translation, the reactions were pelleted through a sucrose cushion to collect ribosomes, and the samples were vitrified on cryo-EM grids. The resulting dataset was processed in RELION 5.0^39^ (Figure S2, Table S1). Initial pre-processing and 3D classification yielded a small subset class of unusually stable collided disomes with well-defined orientations of the stalled and collided ribosomes. Further refinement revealed the presence of elongated density spanning both ribosomes and corresponding to human GCN1 and this subset was further enriched using focused classification with signal subtraction using a mask that protected the GCN1 density from subtraction. We did not observe any density for GCN2 in this class. The low local resolution of the GCN1 density precluded conventional model building. An initial model for human GCN1 was therefore generated in AlphaFold 3^40^ that likely corresponds to an autoinhibited, off-ribosome state (Figure S3A). The model was segmented based on the Alphafold 3 plot of domains of low prediction error (Figure S3B) and adjusted to fit the density roughly (Figure S3C) and improved using PredictandBuild in Phenix^41, 42^, which uses Alphafold 2^43^ to improve and refit the model into the cryo-EM density iteratively (see Methods for details).

As expected^27, 35^, the structure (Figure 1, Figure S4) reveals a stalled ribosome in the canonical state with eRF1^AAQ^ and its binding partner ABCE1 in the A site, a peptidyl tRNA in the P/P state in the P site and a deacylated tRNA in the E site, with the L1 stalk in a closed, locked conformation. The collided ribosome is in the rotated state with A/P and P/E hybrid tRNAs. Density for the 40S of the collided ribosome is well-defined and comparable in resolution to density for the stalled ribosome, but the 60S density of the collided ribosome is less well-defined. The P stalks of both ribosomes are defined well enough to permit rigid body docking of Alphafold 3 models of uL10 and P1/P2 heterodimers. The P stalk of the collided ribosome is stabilized by intimate contacts with the N-terminal region of GCN1. The density is compatible with the Alphafold 3 prediction of this complex (Figure S5).

**Figure 1.**
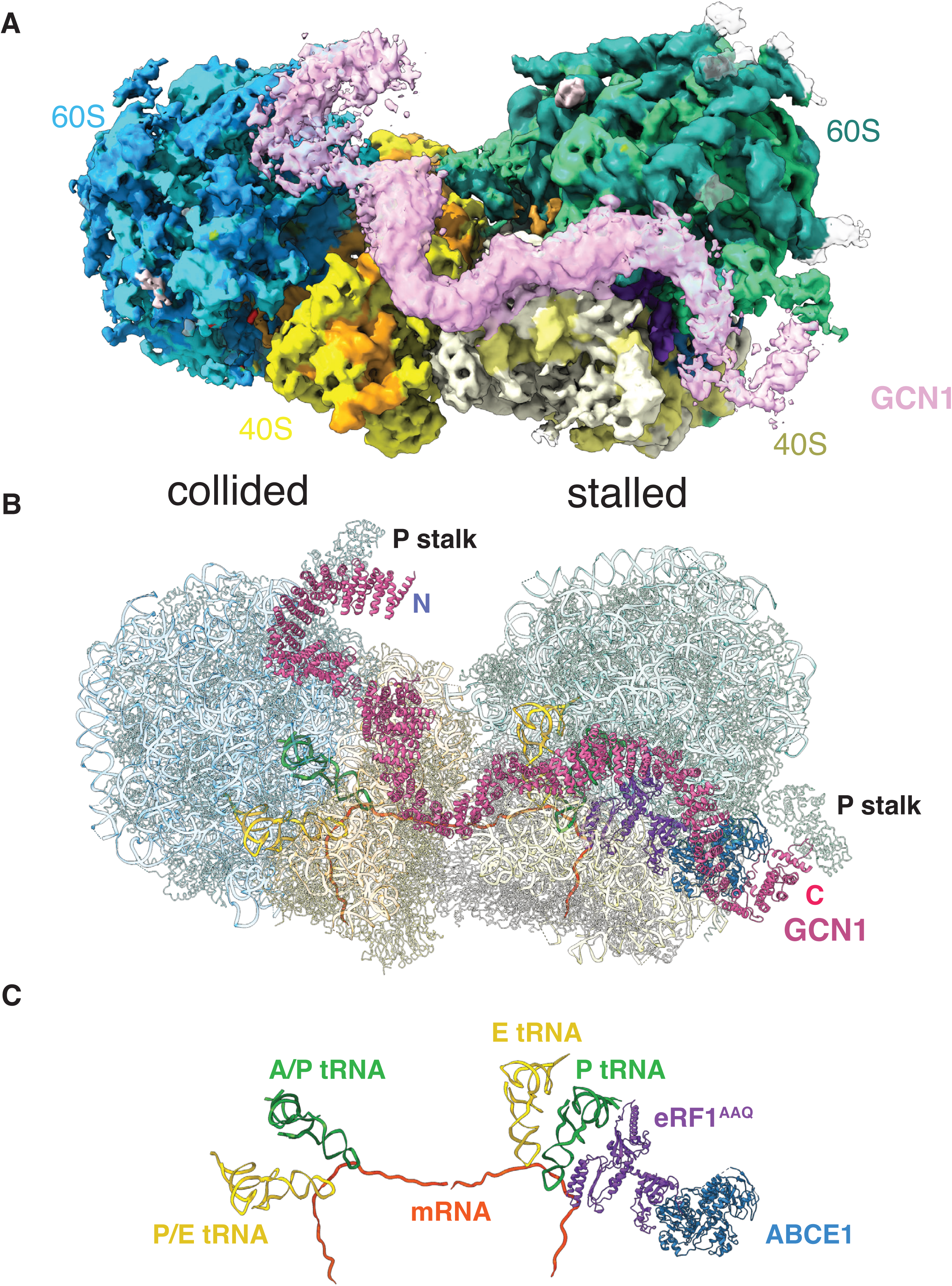
Structure of human GCN1 bound to the rabbit collided di-ribosome. **(A-C)** The stalled ribosome on the right is in the canonical state and is bound to eRF1^AAQ^-ABCE1 in the A site, peptidyl tRNA in the P site and deacylated tRNA in the E site. The collided ribosome on the left is in the rotated state bound to A/P and P/E tRNAs: **(A)** map, **(B)** model and **(C)** bound protein factors and tRNAs. GCN1 spans the P stalk of the collided ribosome (GCN1 N-terminus) to the P stalk of the stalled ribosome (GCN1 C-terminus). To enhance clarity and map continuity, GCN1 density is contoured at a lower threshold than the ribosomes in (A).

### GCN1 binding stabilizes the disome

Our earlier structure of the mammalian collided disome^27^ revealed that the collision interface between ribosomes is flexible and that the bite angle between the stalled and collided ribosomes can vary depending on polysome context (Figure S6). High resolution structure determination of the collided disome therefore required multi-body refinement and principal component analysis of the conformational variability^44^. In contrast, GCN1-bound disomes were sufficiently rigid to obviate multi-body refinement and were instead refined as a single rigid body. This difference is highlighted by a heat-map representation of flexibility in the 40S subunit of the collided ribosome when GCN1 is absent (Figure 2, Unbound versus GCN1-bound). Stable disomes are likely necessary for robust recruitment of GCN2 and the Gcn20-homolog to the collided disome during amino acid starvation to avoid spurious signaling on incidental collisions and on collided disomes that form in other cellular contexts^27, 28, 29, 37, 38^. This result is consistent with recent evidence indicating that GCN1 binding to collided disomes inhibits their ubiquitylation by the RQC pathway^45^.

**Figure 2.**
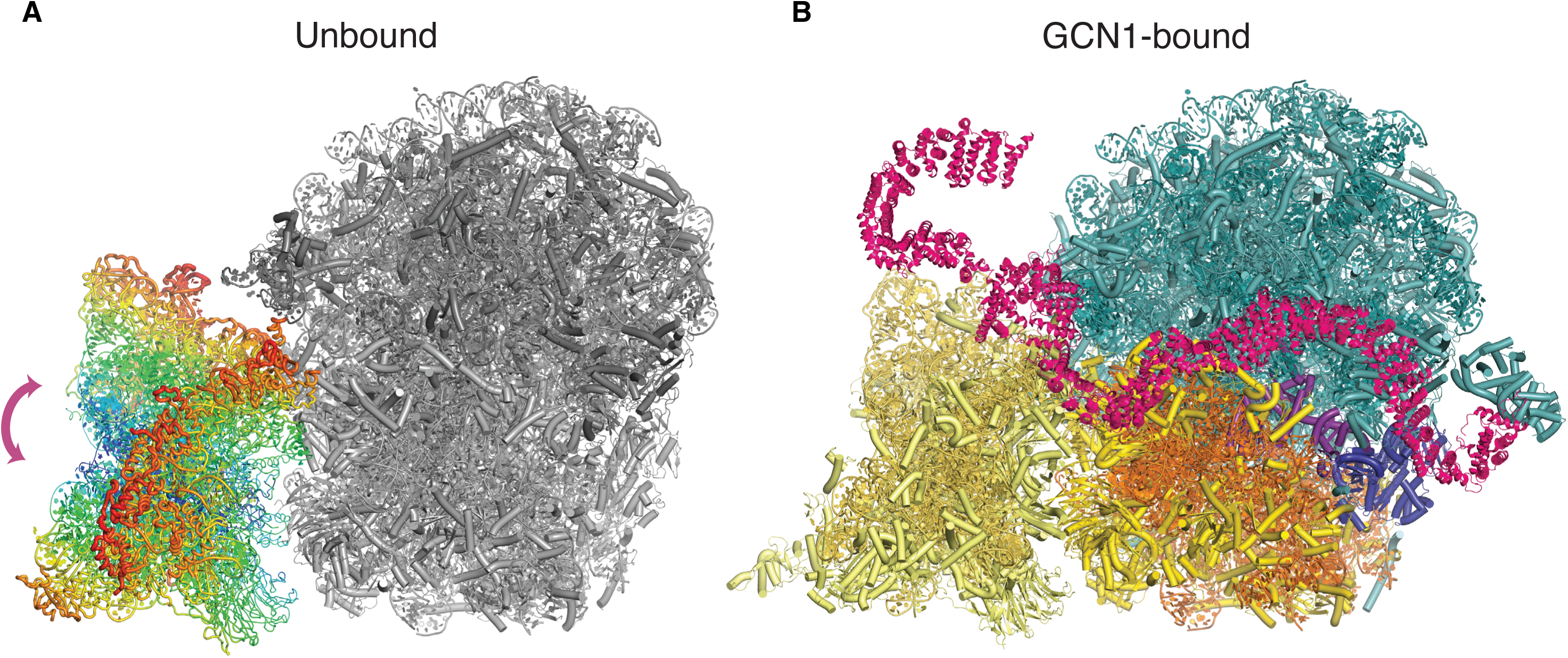
GCN1 stabilizes the collided di-ribosome. **(A-B)** Comparison of the flexibility of the collided di-ribosome (A) with the GCN1-bound collided di-ribosome (B). The flexibility of the 40S of the unbound collided ribosome relative to a static stalled ribosome is quantified by multi-body refinement and principal component analysis and is shown in sausage representation and colored as a heatmap in (A) (mobility decreases from red > orange > yellow > green > blue). This flexibility is absent in the GCN1-bound collided di-ribosome. Also see Figure S2. The 60S of the collided ribosome is hidden for clarity in both (A) and (B). In (B), the 40S proteins are colored yellow, rRNAs – orange, 60S – deep teal, GCN1 – pink, eRF1^AAQ^ – purple, ABCE1 – blue.

### A GCN1 ‘clamp’ contacts the collided ribosome

The N-terminus of GCN1 makes direct contacts with the normally flexible P-stalk of the collided ribosome (Figure 3A). The Alphafold 3 prediction of the P stalk rRNA, protein uL10 and two copies of P1/P2 heterodimers, together with the N-terminal segment of GCN1 (Figure S5) fits well into the observed cryo-EM density in this region (Figure 3A). In addition to this contact, GCN1 also interacts with the beak of the 40S subunit of the collided ribosome via a ‘clamp’ formed by an alpha-helical section and a loop (Figure 3B). Together these elements ‘pinch’ the beak of the collided ribosome presumably to stabilize the interaction. GCN1 also contacts uL5 in the 60S of the stalled ribosome using a loop comprising 1849-1865 and makes extensive contact with the 40S of the stalled ribosome, interacting with uS13 and eS19. Finally, it makes tip-to-tip contact with the P stalk of the stalled ribosome using its C-terminus (Figure 3C). This interaction is likely further stabilized by the presence of GCN2, which is known to interact directly with the P-stalk^13^.

**Figure 3.**
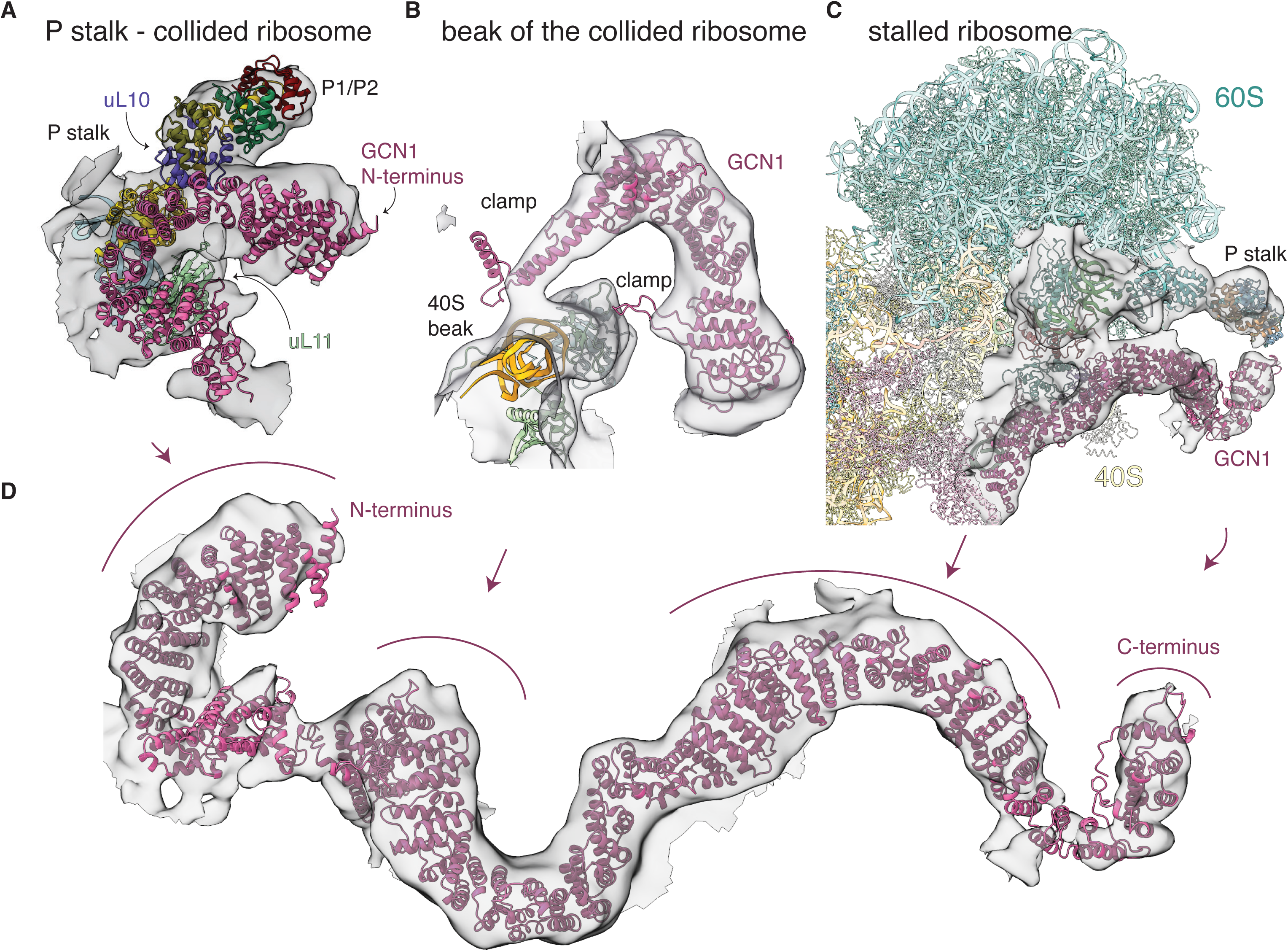
Interactions between GCN1 and the ribosome. **(A)** The N-terminus of GCN1 binds the P-stalk of the collided ribosome, contacting proteins P1, P2, uL10 and uL11. **(B)** ‘Pinching’ interactions between the beak of the collided ribosome and the two components of the clamp region of GCN1. **(C)** Other contacts between GCN1 and the stalled ribosome, including those between the extreme C-terminus of GCN1 and the P stalk. **(D)** overall map-model fit of GCN1. The arrows indicate the positions of the regions depicted in (A)-(C).

### GCN1 - collided ribosome interactions are required for GCN2 activation

To test whether disruption of GCN1 binding to the collided ribosome affects activation of GCN2 *in vivo*, we turned to the yeast system. Alphafold3 modelling revealed that the ‘clamp’ in human GCN1, which interacts with the beak of the 40S subunit of the collided ribosome, is structurally conserved in yeast Gcn1, where the α-helical section and loop correspond to amino acids 746-805 and 1160-1176, respectively (Figure 4A, yellow and red shading). We generated single deletions and a double deletion of these residues in C-terminally Myc-tagged full-length yeast *GCN1* on a low copy plasmid (Figure 4B) and introduced them into a *gcn1Δ* yeast strain. None of these deletions consistently altered steady-state Gcn1-myc protein levels in cell extracts of replicate cultures (Figure S7A). To assess Gcn2 activation, we first tested the strains for sensitivity to 3-aminotriazole (3-AT), which inhibits the histidine biosynthetic enzyme His3 and activates Gcn2, with attendant up-regulation of *GCN4* mRNA translation and transcriptional activation of *HIS3* and many other amino acid biosynthetic genes subject to the general amino acid control. As such, eliminating *GCN2* or *GCN1* impairs growth in medium containing 3-AT^14^. Each single deletion conferred moderate 3-AT-sensitivity (3-AT^S^), whereas the double deletion greatly impaired growth at both 15 mM and 30 mM 3-AT, indistinguishably from the absence of *GCN1* in this strain (Fig. S7C). Consistent with these results, the double deletion, but neither single deletion, significantly diminished the strong induction of Gcn4 protein conferred by 3-AT treatment in all three replicates examined (Figure S7A, Gcn4 & S7B(i), cols. 5-8). None of the mutations, however, significantly decreased the level of eIF2α phosphorylated by Gcn2 on Serine-51, quantified by measuring the ratio of phosphorylated to total eIF2α (Figure S7A, eIF2α-P & S7B(ii), cols. 5-8). (We showed previously that deleting the entire *GCN1* gene abolished the increased eIF2α-P produced under these same starvation conditions, as observed on deleting *GCN2* itself^25^.) Together, these findings suggest that each Gcn1-ribosome contact involving the clamp can independently support nearly WT Gcn2 activation in response to histidine limitation but that eliminating both contacts simultaneously substantially impairs Gcn2-dependent induction of Gcn4 in starved cells.

**Figure 4.**
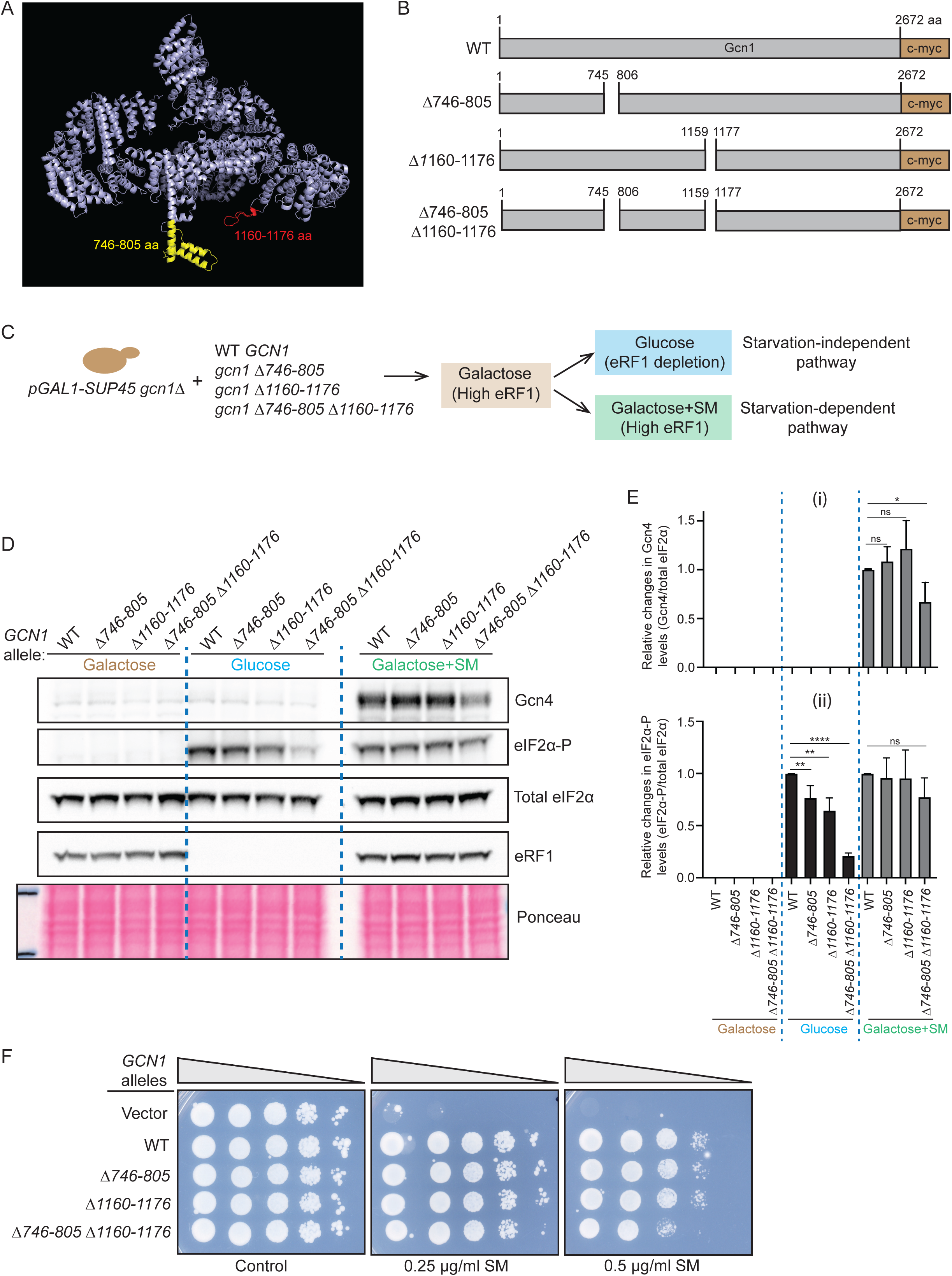
Separate Gcn1 contacts made with the 40S beak of a colliding ribosome function additively to promote full activation of Gcn2 by both starvation-independent and - dependent pathways. **(A)** AlphaFold 3 prediction of yeast Gcn1 structure highlighting the α-helical section and loop comprised of amino acids 746-805 (yellow) and 1160-1176 (red), respectively, corresponding to residues in mammalian GCN1 contacting the 40S beak, which are eliminated by the mutations depicted in panel B. **(B)** Schema of WT Gcn1 protein and variants encoded by *gcn1-Δ746-805*, *gcn1-Δ1160-1176*, and *gcn1-Δ746-805, Δ1160-1176*, all tagged at their C-termini with c-myc epitope. **(C)** Experimental design for transformants of *P_GAL1_-SUP45 gcn1*Δ strain RG3 with the indicated *GCN1* alleles cultured in SC-Ura containing 2% galactose and shifted to SC-Ura containing 2% glucose to deplete eRF1 to analyze the starvation-independent pathway, or shifted to SC-Ura-Ile-Val containing 2% galactose and treated with SM to analyze the starvation-dependent pathway of Gcn2 activation. **(D)** Immunoblot analyses from the experiment outlined in panel C using antibodies against the indicated proteins, with total eIF2α and Ponceau S staining of total proteins providing loading controls. **(E)** Immunoblot signals in (D) for Gcn4 and eIF2α-P were normalized to those for total eIF2α and used with corresponding data from two other biological replicates to calculate the means ± SD shown in the histograms for Gcn4 (i) and eIF2α-P (ii), with asterisks denoting statistical significance determined by a Student’s t-test: *p<0.05, **p<0.01, ****p<0.0001, ns-non-significant difference. **(F)** RG3 transformants with empty vector or the indicated *GCN1-myc* alleles were cultured in SC-Ura containing 2% galactose, serially diluted and spotted on solid SC-Ura-Ile-Val medium containing 2% galactose (control) or to the same medium supplemented with SM at 0.25 μg/ml or 0.5 μg/ml, and incubated for 3-4 days at 30°C.

We went on to examine the effects of these same mutations on Gcn2 activation conferred by collided ribosomes in the absence of amino acid limitation. We showed previously that conditional depletion of translation termination factor eRF1 and attendant stalling of ribosomes at stop codons in the translatome confers Gcn2 activation in a manner requiring at least one of the two P1/P2 heterodimers of the ribosomal P-stalk; whereas neither heterodimer is required for robust Gcn2 activation by 3-AT or the inhibitor of isoleucine/valine biosynthesis sulfometuron methyl (SM). The starvation-independent (SI) pathway of Gcn2 activation is however completely dependent on Gcn1^25^. Depletion of eRF1 was achieved by shifting wild-type or *gcn1-myc* mutant cells expressing eRF1 from a *SUP45* allele driven by the galactose-inducible, glucose-repressible *pGAL1* promoter from galactose-to glucose-containing medium for 12 h (Fig. 4C). As expected, this shift conferred a marked increase in the eIF2α-P/eIF2α ratio in the WT *GCN1-myc* strain (Figure 4D & 4E(ii), cols. 1 & 5). The single deletions of clamp residues conferred moderate, but significant reductions in the phosphorylated fraction of eIF2α, whereas a stronger decline in the eIF2α-P/eIF2α ratio was observed for the double deletion following the shift to glucose (Figure 4D & 4E(ii), cols. 5-8). These findings suggest that the Gcn1 clamp is critically required for activation of Gcn2 by the SI pathway engendered by ribosome stalling without amino acid limitation.

Treating eRF1-replete cells growing on galactose with SM to engage the starvation-dependent pathway also induced eIF2α-P in *GCN1-myc* cells; however, the eIF2α-P/eIF2α ratio was not significantly reduced by any of the three clamp mutations (Figure 4D & 4E(ii), cols. 9-12). In contrast, the double mutation, but neither single clamp deletion, reduced the extent of Gcn4 protein induction conferred by SM treatment in the same cells (Fig. 4D & 4E(i), cols. 9-12); and it also conferred growth sensitivity to two different concentrations of SM (Fig. 4F). These last findings mirror the results obtained in cells starved for histidine with 3-AT shown in Fig. S7 indicating that one or the other ribosome contacts made by the Gcn1 clamp is sufficient for nearly WT Gcn2 activation by the starvation-dependent pathway, while also indicating that considerable activation is retained even without the clamp in cells starved of isoleucine/valine. The fact that the double clamp deletion almost completely eliminated Gcn2 activation in response to eRF1 depletion, while only partially impairing its activation by SM, suggests that the SI pathway has more stringent requirements than the starvation-dependent pathway for proper engagement of Gcn1 with the 40S beak of the colliding ribosome and attendant Gcn2 activation.

The surprising finding that Gcn4 was not induced by eRF1 depletion despite strong induction of eIF2α-P in WT *GCN1-myc* cells (Figure 4D, col. 5) was not completely unexpected considering our previous results using this system, where we observed much stronger induction of Gcn4 by SM versus eRF1 depletion. One possibility is that eRF1 depletion impairs the delayed-reinitiation mechanism of *GCN4* translational control mediated by the four regulatory uORFs, which enables ribosomes to reinitiate at the *GCN4* main ORF after terminating translation at the 5’-proximal uORF1 when levels of the eIF2·GTP·Met-tRNAi ternary complex are reduced by eIF2α phosphorylation in response to amino acid starvation, such as imposed by SM^25^. The failure to observe any Gcn4 induction on eRF1 depletion in the current study might indicate that the plasmid-borne *GCN1-myc* allele employed here does not confer the wild-type level of Gcn1 function produced by the chromosomal *GCN1* allele present in our previous study, further compromising induction of *GCN4* translation.

### ABCF3 is the human ortholog of yeast Gcn20

In yeast, Gcn1 exists as a constitutive heterodimer with Gcn20^19, 30^, an ATPase of unclear function but whose interaction with Gcn1 is required for full Gcn2 activation in amino acid-starved cells^30^. Yeast Gcn20 binds the collided disome near the collision interface^19^. The identity of the human ortholog of yeast Gcn20 is ambiguous. BLAST searches reveal three paralogs as likely candidates – ABCF1, 2 and 3. To determine whether GCN1 interacts at all with a human ortholog of Gcn20, we raised nanobodies against human GCN2 and GCN1 in alpacas (see **Methods** and Figures S8, S9, S10). Human Expi293 cells grown in suspension were treated with halofuginone (HF), known to deprive cells of prolyl-tRNA and activate GCN2^46, 47^ (Figure S11B), and harvested at 0 min, 30 min, 1 h and 18 h post-HF addition. Cells from the 30-min timepoint were lysed, Twin-Strep-tagged GCN2 or GCN1 nanobodies were added to the lysate, and GCN-bound complexes were affinity-enriched and subjected to protein ID mass spectrometry. Gratifyingly, most ribosomal proteins were detected in both samples (Figure S12, Table S2). GCN2 was enriched 29-fold when an anti-GCN2 nanobody was used as bait relative to the sample where an anti-GCN1 nanobody was used as bait. Conversely, GCN1 was enriched 28-fold when a GCN1 nanobody was used. ABCF3 was detected in both samples and was 10-fold more abundant in the anti-GCN1 nanobody sample relative to the anti-GCN2 sample, suggesting a direct interaction between GCN1 and ABCF3 under conditions of HF-mediated depletion of prolyl-tRNA. GCN1 and ABCF3 have indeed previously been suggested to function together in *C. elegans*^48^. However, we cannot yet rule out a role for ABCF1 and 2 in the ISR, which appear to associate more specifically with GCN2.

## Discussion

Here we have demonstrated that ribosome collisions are sufficient to recruit human GCN1. Moreover, GCN1 appears to be agnostic to the cause of the stalling, only requiring the presence of a collided disome in the canonical and rotated states of the stalled and collided ribosomes, respectively. Selectivity for collided disomes that form especially during amino acid starvation may require the presence of other factors, GCN2 for instance. Our cryo-EM results reveal that GCN1 acts as a molecular “splint” that stabilizes the otherwise flexible collision interface between the stalled and collided ribosomes of a disome. This feature may help ensure that GCN2 is only recruited to persistent collisions rather than transient, incidental collisions that can occur during normal translation. Comparison of our structure with the yeast Gcn1-disome complex reveals the overall similarity of the two structures with an RMSD of 3.8 Å between the stalled ribosome-GCN1 complex and 4.2 Å between the GCN1 chains. The differences are due to the constraints of modelling regions of limited resolution, the differences in the factors bound to the yeast and human disomes and the fact that the N-terminus of yeast Gcn1 was absent in the yeast structure. Gratifyingly, our structure is also consistent with an earlier study that visualized the GCN1-collided disome complex *in situ* albeit at low resolution^49^. The direct contact observed here between GCN1 and the P-stalk is significant. Since the P-stalk is a universal recruitment hub for elongation factors (eEF1A/eEF2), GCN1 might physically displace these factors to prioritize stress signaling over continued elongation. The structure also explains why deletions that eliminate the Gcn1 “clamp” result in a “Gcn^-^” (General control nonderepressible) phenotype in yeast, as this element is required for full activation of Gcn2 in amino acid-starved cells (Figures 4 & S7 and ^15^).

Functional analysis of our single and double clamp deletion mutations in yeast Gcn1 suggest that the Gcn1 - collided ribosome interactions are required for WT activation of Gcn2 by both the starvation-dependent and SI pathways *in vivo*. Interestingly, a marked decrease in eIF2α phosphorylation conferred by the double clamp mutation was evident only for the SI pathway, engendered by eRF1 depletion in glucose medium (Figure 4D-E(ii)), whereas for the starvation-dependent pathway, a reduction in Gcn2 activation could only be inferred from the diminished induction of Gcn4 and impaired growth observed in cells starved for amino acids with 3-AT or SM (Figure S7A-C and Figure 4D, 4E(i), and 4F). This comparison may indicate that the functional requirements for Gcn1-mediated activation of Gcn2 are less stringent in starved cells, where uncharged tRNA is present as an activating ligand, than under conditions where stalled ribosomes alone trigger Gcn2 activation.

While yeast Gcn20 and the need for its direct interaction with Gcn1 for robust GCN2 activation *in vivo* has been known for decades^30^, the identity of its human counterpart was obscured by the presence of three paralogs (ABCF1–3). Our nanobody-based affinity purification-mass spectrometry data suggest that ABCF3 is the functional ortholog of Gcn20 found associated with GCN1 during amino acid starvation in mammalian cells. However, both ABCF1 and 2 were also detected in our mass spectrometry data (Table S2), hinting at the possibility of tissue-, or context-specific redundancy in the use of the three paralogs. The involvement of the ABCF ATPase in this pathway is intriguing. It remains to be determined whether the ATPase activity of ABCF3 is required during mammalian ISR activation. Gcn20 and the ABCF proteins are structurally homologous to eEF3, the fungal-specific elongation factor that facilitates evacuation of the E-site tRNA during translation^50^. Recent studies implicate an additional role for eEF3 during termination and post-termination ribosome recycling^51^. Although the exact role of ABCF3’s ATP hydrolysis in GCN2 activation (if any) remains to be fully elucidated, we hypothesize it may regulate the assembly/disassembly of the sensor complex or assist in the eventual rescue of the stalled ribosome once the stress is resolved. Notably, the ATPase domains of yeast Gcn20 enhance but are dispensable for activation of Gcn2, which requires only the Gcn20 N-terminal region that binds Gcn1^30^.

Collision sensors like EDF1, ZNF598 and ZAK also recognize disomes^27, 28, 29, 37, 38^. The observation that GCN1 locks collided disomes into a specific conformation suggests a competitive model where GCN1/2 might dominate during nutrient scarcity to prioritize global translation shutdown (ISR) over mRNA-specific mechanisms such as the RQC^22^. In support of this idea, a recent preprint suggests that GCN1 binding inhibits ubiquitylation of ribosomal proteins by ZNF598 and therefore blocks the RQC pathway^45^. It is possible that GCN1 binding physically locks the 40S subunits into a rigid arrangement that sterically occludes or mechanically prevents the structural rearrangements required for ZNF598 to access and ubiquitylate its target ribosomal proteins. It is also formally possible, however, that different thresholds may trigger ZNF598 versus GCN1 recruitment. For example, a single collision on an mRNA might be handled by ZNF598/RQC to clean up a defective transcript, while more pervasive cell-wide collisions caused by starvation might saturate the RQC system, allowing GCN1 to dominate and activate the ISR. Our model supports the “two-hit” hypothesis: GCN1 recognizes the collided architecture, while GCN2 senses the deacylated tRNA in the A site of the stalled ribosome, consistent with the recent in vitro reconstitution results showing that the presence of cognate uncharged tRNA in the A site leads to higher levels of GCN2 activation than that triggered by ribosome collisions alone^26^. We speculate that the presence of GCN2 would trap any incoming uncharged tRNAs during the decoding of the A-site codon to block accommodation, thus preventing the formation of an unproductive ribosomal state with a fully accommodated, uncharged tRNA in the A site. This intermediate state is an important future target of cryo-EM structure determination and would be reminiscent of the analogous structure of RelA bound to the 70S ribosome during stringent control in bacteria^18^.

The fact that we stalled the lead ribosome with dominant-negative eRF1^AAQ^, which stably occupies the A site, might explain why we did not capture GCN2 on the same collided di-ribosome bound by GCN1, as likely clashes between eRF1 (or the ribosome splitting factor ABCE1 that it recruits) and GCN2 might interfere with GCN2 binding near the A site of the stalled ribosome. Other approaches to induce ribosome collisions, including the use of halofuginone, might facilitate the assembly of GCN1•GCN2 co-complexes for cryo-EM structure determination.

The previously reported yeast Gcn1-disome structure also contained other factors not visualized here, including Rbg2/Gir2 in the A site, eIF5A in the E site and Mbf1 (the yeast homolog of human EDF1) at the collision interface^19^. It is presently unclear whether EDF1 plays a role in recruiting GCN1 binding, perhaps by stabilizing the observed di-ribosome conformation. Our choice to include recombinant EDF1 when assembling the complex was guided by the presence of its yeast ortholog Mbf1 in the structure of yeast Gcn1 bound to disomes^19^. However, we do not see density for EDF1 at the expected location on the 40S-40S collision interface. Future improvements in the resolution of this complex may be necessary to unambiguously answer whether EDF1 is present on the GCN1 di-ribosome complex. Interestingly, however, we did detect the mammalian ortholog of yeast Rbg2, namely DRG2, in our nanobody proteomic dataset (Table S2) and future studies will focus on elucidating the status and contribution of the stalled ribosomal A site in atomic detail.

The GCN1 family also plays a role in surveillance mechanisms that respond to stalled ribosomes with an occluded A site^52^. In this case, the E3 ubiquitin ligases RNF14 and RNF25 are recruited to ubiquitylate eukaryotic elongation factor 1A (eEF1A) and specific ribosomal proteins. It is presently unclear how this role of GCN1 is related to GCN2 activation during amino acid starvation. Another intriguing role has also been assigned to GCN1 in co-translational degradation of mRNAs with translation readthrough into the 3’ UTR, in concert with the deadenylase CCR4-NOT^53^.

The conservation of the “clamp” region from yeast to humans underscores the fundamental nature of this architecture. However, the apparent differences in affinity of Gcn1 for constitutively-bound Gcn20 in yeast versus GCN1 and more weakly-associating ABCF3 in humans is reminiscent of the different affinities for the E3 ubiquitin ligase subunit Not4 (CNOT4 in human) in another ribosome stalling sensor, CCR4-NOT^54, 55^ and might reflect an additional layer of regulation in higher eukaryotes that integrates inputs from other metabolic pathways or eIF2*a* kinases besides GCN2.

## Methods

### Protein Constructs

pFastBac1-His_6_-Twin-Strep-tag-TEV-GCN1 and GCN2 plasmids were kind gifts from Roger Williams^13^. Baculoviruses were generated according to the Bac-to-bac manual (Life Technologies) using DH10EMBacY cells and transfected in suspension into Sf9 insect cells grown in Sf-900 II SFM media using either Fugene HD or Cellfectin II transfection reagents. P0, P1 and P2 viruses were generated and the P2 virus stock was used to infect 4 L of Hi5 cells at an MoI of 5 and the cells were incubated with shaking at 140 rpm at 27 °C for 51-54 hours post infection. Cells were harvested by centrifugation at 2000 x g for 15 minutes and the cell pellets were stored at – 80 °C until use. Human EDF1 (residues 2-148) was cloned with an N-terminal His_6_-SUMO tag into pCA528 (AddGene #191234) for *E. coli* expression.

#### GCN2

Cells were lysed in binding buffer (50 mM TRIS pH 8, 150 mM NaCl, 5% glycerol, 0.5 mM TCEP) supplemented with 1.5% TRITON X-100 + 1 mM PMSF + cOmplete EDTA-free protease inhibitor cocktail, the resuspension was sonicated and clarified by centrifugation in a JA-25.50 rotor (Beckman Coulter) for 30 minutes at 20000 rpm and at 4 °C. The clarified supernatant was filtered using 0.45 µm syringe filters and loaded onto a 5 ml StrepTrap XT column at 3 ml/min. The column was washed with 10 column volumes (CV) binding buffer supplemented with 1 M NaCl, followed by 10 CV of binding buffer and eluted with 1x buffer BXT (IBA biosciences) supplemented with 5% glycerol and 1 mM TCEP pH 7. The eluate was concentrated down to 2 ml using 50 kDa molecular weight cutoff concentrators and subjected to size exclusion chromatography on a Superose 6 column (Cytiva) at 1 ml/min in 50 mM HEPES pH 7.5, 100 mM KOAc, 5% glycerol, 1 mM TCEP. Fractions corresponding to the protein peak were pooled and the protein concentrated to 105 µM, aliquoted, flash-frozen in liquid nitrogen and stored at – 80 °C.

#### GCN1

Cells were lysed in binding buffer (50 mM TRIS pH 8, 150 mM NaCl, 5% glycerol, 1 mM DTT) supplemented with cOmplete EDTA-free protease inhibitor cocktail and bio-lock (IBA), the resuspension was lysed using 30 strokes in a 100 ml dounce homogenizer and clarified by centrifugation in a JA-25.50 rotor (Beckman Coulter) for 20 minutes at 20000 rpm and at 4 °C. After filtering the supernatant through 5 µm syringe filters, the supernatant was sonicated for 2 minutes at 70% power at 4 °C and filtered through 0.5 µm syringe filters. The filtered supernatant was loaded onto a 5 ml StrepTrap XT column at 3 ml/min, the column washed with 10 CV binding buffer supplemented with 1 M NaCl, followed by 10 CV binding buffer and the protein eluted with 1x buffer BXT + 5 % glycerol + 1 mM DTT. The pooled fractions were subjected to Q anion exchange chromatography using 2 x 5 ml HiTrap Q HP columns (Cytiva) connected in series and eluted in a 200 ml gradient from 150 mM NaCl to 1 M NaCl in binding buffer. Pooled fractions were concentrated down to 3 ml and loaded onto a Superdex 200 size exclusion column equilibrated in 50 mM HEPES pH 7.5, 200 mM NaCl, 5% glycerol, 1 mM DTT. Pooled fractions of GCN1 were concentrated to 80 µM, aliquoted, flash-frozen in liquid nitrogen and stored at – 80 °C.

#### EDF1

pCA528-His_6_-SUMO-EDF1 was transformed into *E. coli* BL21 (DE3) and 2 L cultured overnight at 37 °C in terrific broth-based autoinduction medium with 0.5% glycerol, 0.2% lactose and 0.05% glucose. The cells were harvested, resuspended in 1x PBS supplemented with NaCl to a final concentration of 300 mM and stored at -80 °C. The cells were thawed, cOmplete EDTA-free protease inhibitor cocktail added and TRITON X-100 was added to a final concentration of 1.5%. Cell lysis was aided by addition of 0.5 mg lysozyme, 4 mM MgCl_2_, 4 mM CaCl_2_ and 0.5 mg DNAse I. Following a 20-minute stirred incubation at 23 °C, the lysate was sonicated on ice for 10 minutes at 50% power with a 10s on-off pulse. The lysate was centrifuged in a Beckman JA-25.50 rotor at 21000 rpm for 15 minutes at 4 °C and the clarified supernatant filtered through 0.45 µm syringe filters. The filtered supernatant was incubated with 5 ml of Ni-NTA resin (Qiagen) equilibrated in binding buffer (100 mM Na Phosphate pH 7.4, 300 mM NaCl) for 1 hour at 4 °C with rocking, the flowthrough was removed and the drained beads washed with 250 ml of binding buffer, 250 ml of binding buffer with 1 M NaCl and 250 ml of binding buffer again. The bound proteins were eluted with 10 fractions of 5 ml each of binding buffer supplemented with 250 mM imidazole and pooled. 225 µg of home-made Ulp1 was added and the mixture was dialyzed into 1x PBS (with 300 mM NaCl) + 0.1 mM TCEP. After verification of complete tag cleavage by SDS-PAGE, the Ulp1 and cleaved His_6_-SUMO tag were removed by rebinding to Ni-NTA beads. The flowthrough comprised pure, untagged EDF1 and was concentrated to 3.4 mM (55.4 mg/ml), aliquoted, flash-frozen and stored at – 80 °C.

#### GCN2 and GCN1 nanobodies

Twin-Strep-tagged versions of the nanobodies (see section below) were cloned into pRSETA vectors, expressed in *E. coli* C3026 Shuffle-T7 cells (NEB), cultured in a terrific broth-based autoinduction medium at 19 °C overnight and purified via Strep-Tactin XT affinity chromatography, Q anion and SP cation exchange chromatography, concentrated to 3.8 mg/ml, flash-frozen in liquid nitrogen and stored at – 80 °C until use.

### In vitro translation

Translations were performed at 32 °C using nuclease-untreated rabbit reticulocyte lysates (RRL) in the presence of ^35^S-L-methionine (10.25 µCi; Revvity) or 40 µM cold L-methionine. Radiolabeling assays were performed at a 20 µl scale and cold reactions were 1 ml in volume. Stalling was induced by including 2 µM eRF1^AAQ^ in the reactions in the presence of 10 µM EDF1. Reactions were pelleted through a 20% sucrose cushion in 1x RNC buffer (50 mM HEPES pH 7.5, 100 mM KOAc, 5 mM Mg(OAc)_2_, 1 mM DTT) and resuspended in 50 µl of 1x RNC buffer lacking Mg^2+^. The ribosome concentration was measured by A_260 nm_ to be 820 nM.

### Complex assembly and sample preparation for cryo-EM

Complex assembly was performed with 205 nM eRF1^AAQ^-stalled polysomes, 1.5 µM EDF1, 0.5 mM AMPPCP, 1 µM GCN2, 0.1 - 0.5 µM GCN1 and adjusted to a final volume of 100 µl using 1x RNC + 1 mM DTT. 3 µl of the sample was vitrified using a Vitrobot Mark IV (Thermo Scientific) onto UltraFoil R3.5/1 grids using a blot force of -15, blot time of 5 s at 4 °C and 100% relative humidity and plunging into liquid ethane held at 91 K. Grids were stored in liquid nitrogen until use.

### Data Collection

34,040 Tiffs (5760 pixel x 4092 pixel x 55 frames) were collected on a 300 kV Titan Krios G3 (Thermo Scientific) equipped with an X-FEG and a K3 camera in super-resolution (bin 2) counting mode and using EPU software by performing aberration-free image shift (AFIS). The pixel size used was 0.826 Åpixel^-1^ and the flux used was 23.3 e^-^Å^-2^s^-1^ for 2 seconds of exposure to yield a total dose of 46.6 e^-^Å^-2^, such that the dose per frame was 0.847 e^-^Å^-2^ frame^-1^.

### Data Processing

Data were processed in RELION 5.0^39^. Movies were aligned using 5 x 5 patches and dose-weighting in RELION’s implementation of MOTIONCOR 2^56^. Contrast transfer function (CTF) was estimated using CTFFIND-4.1^57^ and 30,868 good micrographs corresponding to a CTF maximum resolution less than 7 Å and a CTF figure of merit greater than 0.08 were selected. 904,032 particles were picked using Topaz and downscaled to a 100-pixel box corresponding to 4.13 Å pixel^-1^. An initial 3D reference derived from EMD-4133 was used to perform masked 3D classification with alignments on the picked particles into 10 classes and limiting the maximum number of significant samples (_rlnNrOfSignificantSamples) using (-- maxsig 3000). 549,918 particles corresponding to good ribosomes were selected for further processing, 3D refined to an angular accuracy of 0.986° and a final resolution of 8.4 Å. These particles were reextracted into a 360-pixel box corresponding to a pixel size of 1.147 Åpixel^-1^ and re-refined to an angular accuracy of 0.451° and a resolution of 3.56 Å. These particles were 3D classified without alignments into 15 classes and a class corresponding to 9,432 particles that appeared to comprise unusually stable di-ribosomes was selected and 3D refined to an angular accuracy of 0.258° and a resolution of 5.1 Å. The particles in this class were re-centered on the interface between the stalled and collided ribosomes and re-extracted in a 360-pixel box corresponding to a pixel size of 2.294 Åpixel^-1^. These di-ribosomes were refined to an angular accuracy of 0.42° and 8.26 Å resolution. Maps of the stalled and collided ribosomes from our earlier structure of the collided di-ribosome^27^ were docked into the collided di-ribosome map to create a soft mask for further processing. Putative density for human GCN1 was masked and these particles subjected to focused classification with partial signal subtraction (FCwSS) against this mask using various **τ**2_fudge (T) values. The successful classification used a T of 38 and 10 classes without alignments. A class comprising 1,820 particles with clear GCN1 density was used for final refinement to 10.1 Å local resolution estimation, postprocessing and modelling.

### Model Building and Real Space Refinement

PDB 6CHM and 6CHQ corresponding to the stalled and collided ribosomes^27^ were docked into the density in UCSF Chimera^58^. An Alphafold 3 model for human GCN1 was divided into 5 segments and each piece was docked manually into the GCN1 density to yield a ‘scaffold model’. GCN1 primary sequence and the scaffold model were provided as inputs to Phenix PredictAndBuild^41, 42^. Briefly, models were predicted using AlphaFold2 and the predicted models were trimmed to remove low confidence regions and split into compact sections. The trimmed domains were sequentially docked from all the predicted models into the map and rearranged to connect sequential parts. The rearranged docked model was used as the scaffold model for the next iteration. PredictAndBuild then iterated model prediction using chains from the rebuilt model as templates to improve model prediction compared to a single prediction step. The entire procedure was repeated until the change in predicted models between subsequent cycles was small. The entire model was real space refined against the map in phenix RealSpaceRefine using defaults but with a non-bonded weight of 1000. Rotamers were fit using ‘outliers_or_poormap’ with a sigma of 5.0 and a target of ‘fix_outliers’ and a tuneup of ‘outliers_and_poormap’.

### Yeast strains and plasmids

All yeast strains and plasmids used in this study are listed in Tables S3 and S4, respectively. Plasmids RG20, RG21, and RG22 were synthesized by Life Science Biotechnology using p2367 as the template plasmid.

### Media and growth conditions

Transformants of *gcn1Δ* strain H2079 harboring different plasmid-borne *GCN1-myc* alleles were grown overnight at 30°C in synthetic complete medium lacking uracil (SC-Ura), comprised of 0.5% ammonium sulfate, 0.14% yeast nitrogen base without amino acids and ammonium sulfate, 2% glucose and supplements of all other amino acids. Cells were sub-cultured in the same medium (control addition) or in SC-Ura also lacking histidine (SC-Ura-His) (for 3-AT treatment) and grown until exponential phase (OD_600_ of 0.8), after which cells were harvested (control condition) or incubated for 1h after adding 3-AT to 10 mM. Transformants of *P_GAL1_-SUP45 gcn1Δ* strain RG3 harboring different plasmid-borne *GCN1-myc* alleles were grown overnight at 30°C in SC-Ura containing 2% galactose in place of glucose, sub-cultured in the same medium and grown to exponential phase (OD_600_ of 0.8), washed three times with SC-Ura containing 2% glucose medium and transferred to the same medium and incubated for 12h. Alternatively, cells were sub-cultured in galactose-containing SC medium lacking uracil, isoleucine and valine (SC-Ura-Ile-Val) to OD_600_ of 0.8, SM was added at 0.5 μg/ml, and incubation was continued for 1h.

### Immunoblot analysis

To quantify Gcn1-myc expression, harvested cells were lysed by vortexing with glass beads in lysis buffer (20 mM Tris-HCl [pH 7.5], 50 mM KCl, 10 mM MgCl2, 1 mM DTT) supplemented with protease inhibitors. Protein concentrations of the resulting whole cell extracts (WCEs) were determined using a Pierce bicinchoninic acid (BCA) assay kit (Thermo Scientific, #23227). Total proteins were separated by SDS-PAGE on 4-20% TGX gels (Bio-rad, #5671093). Western immunoblotting was performed with 1:1000 dilution of anti-Myc mouse antibodies (Roche, #11667203001) and immune complexes were detected using HRP-conjugated secondary anti-mouse antibodies (Amersham, #NA931V) and enhanced chemiluminescence (ECL) reagent (Thermo Scientific, #34580). To quantify expression of Gcn4 or phosphorylated eIF2α, cells were lysed using trichloroacetic acid extraction^59^, and protein concentrations determined as above. Total proteins were separated by SDS-PAGE on 10% NuPAGE Bis-Tris gels (Invitrogen, #WG1202). Western blotting was performed with 1:1000 dilution of anti-phospho-eIF2α (Ser51) (Cell Signaling Technology, #9721S), 1:1000 dilution of anti-total eIF2α (CM-217, gift from Thomas Dever), 1:2000 dilution of anti-Gcn4^60^, or 1:1000 dilution of anti-eRF1^61^ antibodies, using HRP-conjugated secondary anti-rabbit antibodies (Amersham, #NA9340V) and ECL as above. All Western signals were quantified using NIH ImageJ software and plotted using Prism 10 software.

### Spotting assays of yeast cell growth

Yeast cells were grown to exponential phase in SC-Ura containing 2% galactose medium for RG3 transformants and in SC-Ura containing 2% glucose for H2079 transformants, harvested by centrifugation, and then washed once, resuspended, and diluted in sterile water. Aliquots of 10 μL of each suspension or dilution were spotted on solid medium and incubated at 30°C for 3-4 days.

### GCN1 and GCN2 Nanobody Generation and Selection

An alpaca (*Vicugna pacos*) was immunized with purified full-length recombinant GCN1 and GCN2 proteins following established nanobody generation protocols^62, 63^. The animal received four subcutaneous immunizations at two-week intervals using antigen formulated with GERBU adjuvant. Serum samples collected during the immunization campaign showed a robust antigen-specific immune response against both GCN1 and GCN2 antigens as determined by ELISA^63^ (Figure S8).

#### Immune Alpaca serum ELISA

Thermo Scientific™ Pierce™ Streptavidin coated plates were incubated with biotinylated recombinant GCN1 or GCN2 proteins (100 nM) for 1 h at room temperature. Control wells received biotinylated chicken lysozyme protein (GeneTex, Cat. No. GTX82960-pro). Following blocking and washing steps, alpaca serum samples were diluted 1:200 and 1:800 in PBS and incubated in antigen-coated wells. Bound antibodies were detected using Peroxidase AffiniPure® Goat Anti-Alpaca IgG (H+L) (Jackson ImmunoResearch, Cat. No. 128-035-003). Plates were developed with 100 μL 3,3′,5,5′-Tetramethylbenzidine (TMB) substrate, and the reaction was stopped with an equal volume of ELISA stop solution (Invitrogen). Absorbance of the TMB reaction product was measured at 450 nm. ELISA signals were compared between GCN1/GCN2 antigen-coated wells and the lysozyme control to assess antigen-specific binding.

Peripheral blood mononuclear cells were isolated following the final boost, and total RNA was extracted for cDNA synthesis. VHH-encoding sequences were amplified by nested PCR and cloned into a phage-display vector using FX cloning^64, 65^. Transformation into Escherichia coli generated a nanobody library containing approximately 10⁸ independent transformants.

#### Antigen biotinylation

GCN1 at 3.6 µM and GCN2 at 117 µM were biotinylated by addition of a 40-fold molar excess of EZ-Link NHS-PEG4-biotin (Thermo Fisher Scientific) followed by incubation at 23 °C for 30 minutes. Excess reagent was removed by dialysis for 2 hours at 23 °C into 2 changes of 1 L of 50 mM HEPES pH 7.5, 200 mM NaCl, 0.25 mM TCEP using 10 kDa molecular weight cutoff Slide-A-Lyzer MINI dialysis devices (Thermo Scientific). Nanobody libraries were subjected to two rounds of phage-display selection against biotinylated GCN1 and GCN2 immobilized on streptavidin-coated magnetic beads. Enrichment of antigen-specific binders was observed following iterative selection, after which the enriched libraries were subcloned into a bacterial expression vector for soluble nanobody production. Individual clones were screened by ELISA against the target antigens, resulting in the identification of multiple GCN1 and GCN2 binding nanobodies (Figure S9). Sequence analysis revealed 8 distinct nanobody families. Representative nanobodies were expressed in E. coli and purified as previously described^66^.

### Fluorescence Anisotropy

Binding interactions with GCN1 and GCN2 were evaluated using fluorescence polarization assays employing Alexa Fluor 488 labelled nanobodies. Purified nanobodies were labelled with Alexa Fluor 488 C5 maleimide (Thermo Fisher Scientific) according to the manufacturer’s instructions using a 10-fold molar excess of dye. Briefly, 100 ul of purified nanobody was incubated with the fluorophore for 1 hour at room temperature and excess dye was removed by desalting before use.

Fluorescent polarisation spectroscopy measurements were subsequently performed using the labelled nanobodies to determine binding affinities.

All reactions were carried out in 20 mM HEPES (pH 7.4), 150 mM NaCl and 0.01% Tween-20. A three-fold serial dilution of GCN2 was prepared, with a final starting concentration of 325 nM following 1:1 mixing with 5 nM Alexa Fluor 488 maleimide-labelled GCN2 nanobody 48. For GCN1, a three-fold serial dilution was prepared, with a final starting concentration of 100 nM following 1:1 mixing with 5 nM Alexa Fluor 488-labelled nanobody. Reactions were carried out in a total volume of 60 µL in a black, flat-bottom, non-binding surface 384-well plate (Corning) at 25 °C. Fluorescence anisotropy measurements were performed using a PheraSTAR FSX plate reader (BMG Labtech) using an optic module for λ_ex_ = 485 nm and λ_em_ = 520 nm, after initial mixing or following a 20 min incubation at room temperature. Data were analysed in PRISM 10 (GraphPad Software) using a single-site ligand-depletion model.

Several nanobodies exhibited high-affinity binding, with apparent dissociation constants ranging from sub-nanomolar to low nanomolar values (Figure S10). The highest-affinity binders were selected for subsequent use.

### Human Cell Culture and GCN1/2 affinity enrichment

Expi293 cells (Thermo Scientific) were seeded at 0.35 x 10^6^ cells ml^-1^ in Expi293 medium and grown overnight to 1 x 10^6^ cells ml^-1^ at 37 °C, 8% CO_2_. 10 mM halofuginone (HF) dissolved in DMSO was added to a final concentration of 200 nM. 2 x 50 ml of cells were pelleted at 500 x g and frozen at -80 °C either before HF addition (i.e., T=0) or after 30 min, 1 hour or 18 hours post-HF addition. 50 ml of cells were lysed in 1 ml of mammalian lysis buffer (50 mM HEPES pH 7.5, 100 mM potassium acetate, 0.5 mM magnesium acetate, 0.01% digitonin, cOmplete EDTA-free protease inhibitor cocktail (Roche), bio-lock (IBA biosciences), 2 mM DTT and flash-frozen in liquid nitrogen. The resuspensions were thawed at 23 °C and lysed by passing through a 26-gauge needle 5 times. The lysate was clarified by centrifugation at 16,100 x g for 8 minutes and the supernatant was saved. An affinity pulldown was performed in which 50 µl of Twin-Strep-tagged GCN2 nanobody or 10 µl of Twin-Strep-tagged GCN1 nanobody was directly added to 1 ml of the supernatant and incubated with Strep Tactin 4-flow resin (IBA biosciences) for 20 minutes at 4 °C with tumbling. The beads were washed once with mammalian lysis buffer followed by 4 washes with RNC-pellet buffer (50 mM HEPES pH 7.5, 100 mM potassium acetate, 0.5 mM magnesium acetate and 0.5 mM TCEP). Affinity-enriched species were eluted in 40 µl of 1x buffer BXT (IBA biosciences) supplemented with RNC-pellet buffer and analyzed by SDS-PAGE.

### Nanobody Proteomics

The SDS-PAGE gels were stained with InstantBlue (Abcam), fragmented using a scalpel and suspended in 100 µl ultrapure water. The anti-GCN2 nanobody used was AGSSSQVQLVESGGGLVPAGGSLRLSCAASGLTFSRYAMGWFRQAPGKEREFVAGIGWRSDANTYYTDSVKGRFTISRDNAKNTVFLQMISLKPEDTATYYCAAEALGDGAGYRRYEYWSKGTPVTVSAGRAGEQKLISEEDLNSAVD. The anti-GCN1 nanobody used was AGSSSQRQLVESGGGLVQPGGSLRLSCAASRIIFAVDRMGMGWYRQPPGKQREFVAHITRGGSTNYADSVKGRFTISRDNDKNTMYLQMNSLKPEDTAVYYCLMRAGASDYWGKGTPVTVSAGRAGEQKLISEEDLNSAVD.

#### In-gel digestion

Gel samples were de-stained with 50% v/v acetonitrile and 200µl of 50 mM ammonium bicarbonate was added and gently mixed for 10 minutes. Proteins were reduced with 10 mM DTT diluted in 100 mM Ammonium bicarbonate for 20 minutes at 55 °C. Solvent was removed and the gels were cooled before alkylation with 55 mM iodoacetamide diluted in 100 mM ammonium bicarbonate. Samples were placed in the dark at room temperature for 30 minutes. Solvent was removed and the gels were washed with 100 mM ammonium bicarbonate. Gel samples were washed with acetonitrile and dried. Samples were then digested with 25 µl of trypsin (Promega, UK) at 6 ng/μl overnight at 37 °C. Peptides were extracted in 2% v/v formic acid, 3% v/v acetonitrile and de-salted.

#### C18 Peptide Clean-up Protocol

A C18 de-salting tip made by adding a plug of C18 material to a 200 µl tip. The tip was wetted and equilibrated by adding 50 μL of Solution B (50% water + 50% acetonitrile (ACN) + 0.1% formic acid (FA)) and spun at 3500 rpm for 2 minutes and this process was repeated once. Then 50 µl of Solution A (100% water + 0.1% FA) was added and the tips spun at 3500 rpm for 2 minutes. This wash was repeated until 3 total washes were performed. 25 μL of sample was applied and bound by slowly spinning at 2000 rpm for 3 minutes and this step was repeated twice more. To de-salt, 50 μL of Solution A was added and removed by spinning at 3500 rpm for 2 minutes. This wash was repeated thrice more. The sample was eluted with 50 μL of Solution C (25% water + 70% ACN + 0.1% FA) by spinning at 3000 rpm and repeated so that the final eluate was in a volume of 100 µl. A SpeedVac was used to remove the acetonitrile and the sample was made up to 30 µl with 0.1% FA and 3% ACN. 6 µl was injected per sample.

#### LC-MS/MS analysis

The extracted tryptic peptides were analysed by nano-scale capillary LC-MS/MS using an Ultimate U3000 HPLC (ThermoFisher Scientific) to deliver a flow of approximately 300 nL/min. The sample was loaded onto the trapping column (Thermo Scientific, PepMap100, C18, 300 μm X 5 mm) and resolved on the analytical column (Aurora 25cm column) at a flow rate of 300 nL/min using a gradient of 100% A (0.1% formic acid) 0% B (80% acetonitrile 0.1% formic acid) to 40% B over 40 minutes. Peptides were analysed, with a hybrid linear quadrupole ion trap mass spectrometer (Orbitrap QE PLUS, ThermoScientific). Data dependent analysis was carried out, using a resolution of 70,000 for the full MS spectrum, followed by ten MS/MS spectra in the linear ion trap. MS spectra were collected over a m/z range of 200–1800. MS/MS scans were collected using threshold energy of 35 for collision induced dissociation.

#### Bioinformatic analysis

Raw data were imported and data processed in Proteome Discoverer v3.1 (Thermo Fisher Scientific). The raw files were submitted to a database search using Proteome Discoverer with SequestHF against the Uniprot reference proteome for human including the protein sequences supplied. Processing step consisted a double iterative search using inferis rescoring on a first pass without any additional protein modifications and spectra with a confidence filter worse than “high” were researched with Sequest HT including additional common protein modifications (Deamidated N,Q; Oxidation M, gln to pyro-Glu Q; N-terminal acetylation; N-terminal acetylation and Methionine loss; Methionine loss. The spectra identification was performed with the following parameters: MS accuracy, 10 ppm; MS/MS accuracy of 0.02 Da for spectra acquired in Orbitrap mass analyzer; up to two missed cleavage sites allowed; carbamidomethylation of cysteine; and oxidation of methionine as variable modifications. Percolator node was used for false discovery rate estimation and only rank 1 peptide identifications of high confidence (FDR<1%) were accepted.

#### Proteomic Data Filtering and Standardization

Raw protein datasets were analyzed via label-free quantitative proteomics using both Proteome Discoverer and Scaffold data outputs to compare differential enrichment between the anti-GCN1 and anti-GCN2 nanobody immunoaffinity purifications in human Expi293 cells under conditions of halofuginone-mediated inhibition of prolyl-tRNA synthesis. To ensure analytical robustness and eliminate artifacts generated by single-sample stochastic detection, any protein missing a quantitative value or reporting an abundance value of zero in either of the two experimental arms was excluded. This strict filtration guaranteed that the analysis was restricted exclusively to the shared interactome quantified concurrently across both nanobody pulldowns. To visualize the comparative preference of the shared translation machinery for either bait, the filtered abundance data were subjected to a coordinate rotation mapping the dataset onto a modified Bland-Altman symmetry plot. A baseline correction factor of +1 was applied to both raw abundance values to prevent mathematical division-by-zero and logarithmic errors. The relative enrichment preference (X-axis) was calculated as the log_2_ fold change of the normalized abundance ratios between the two samples, defined as:

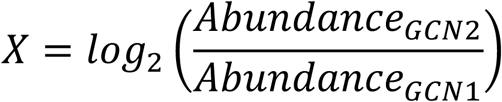

where a vertical baseline spine is established at X=0 for shared proteins captured with equal efficiency. Left-shifted negative values denote preferential enrichment on the GCN2-interactor side, while right-shifted positive values denote a preference for the GCN1-interactor side. The absolute scale of total protein abundance (Y-axis) was represented using a logarithmic baseline of the mean abundance between both captures, defined as:

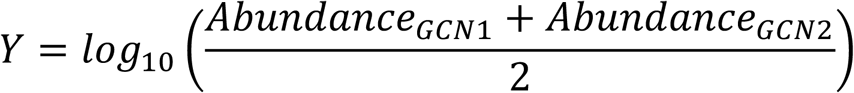

Proteins were computationally stratified into distinct categorical data series based on their annotated descriptions and accession numbers—including Primary Baits (GCN1 and GCN2), ABCF family members, core ribosomal proteins, and general translation factors—allowing background proteins to be rendered with transparency to visually isolate the vertical ribosomal spine. The final interactive visualization was compiled in Microsoft Excel for Mac (v16.109) and exported via object-level vector formatting directly into Adobe Illustrator.

## Acknowledgements

We thank the MRC Laboratory of Molecular Biology (LMB) Electron Microscopy Facility for assistance with cryo-EM data collection and the Scientific Computing Facility for support with high-performance computational resources and data processing. Mass spectrometry analysis was performed at the Biological Mass Spectrometry and Proteomics Facility of the Medical Research Council Laboratory of Molecular Biology, Cambridge, UK. The authors would like to thank Dr Catarina Franco and Dr Farida Begum for the help with sample preparation and data analysis. We thank Debra Davidson for carrying out Alphafold 3 predictions. Coordinates and maps are available from the PDB (36LR) and EMDB (EMD-77635) databases respectively. Raw micrographs have been uploaded to EMPIAR (EMPIAR-13710). This work was supported by the UK Medical Research Council MC_U105184332, a Wellcome Trust Senior Investigator award (WT096570), the Agouron Institute, and the Louis-Jeantet Foundation to VR.

---

**Table S1.**
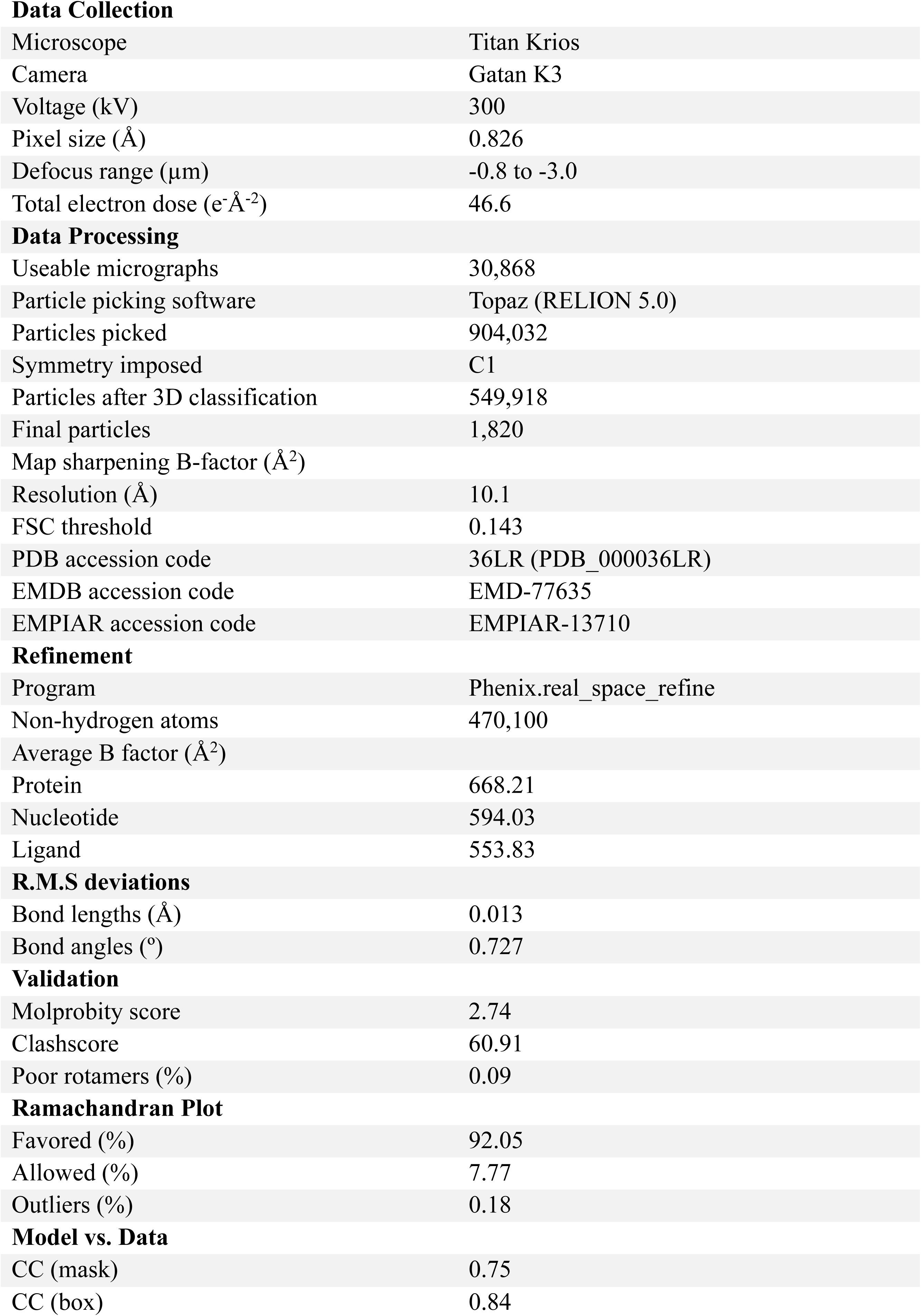
Data collection, processing, refinement and model statistics.

**Table S2 | Nanobody Proteomics**

See attached Microsoft Excel file (Chandrasekaran_et_al_2026_Table_S2.xlsx)

**Table S3.**
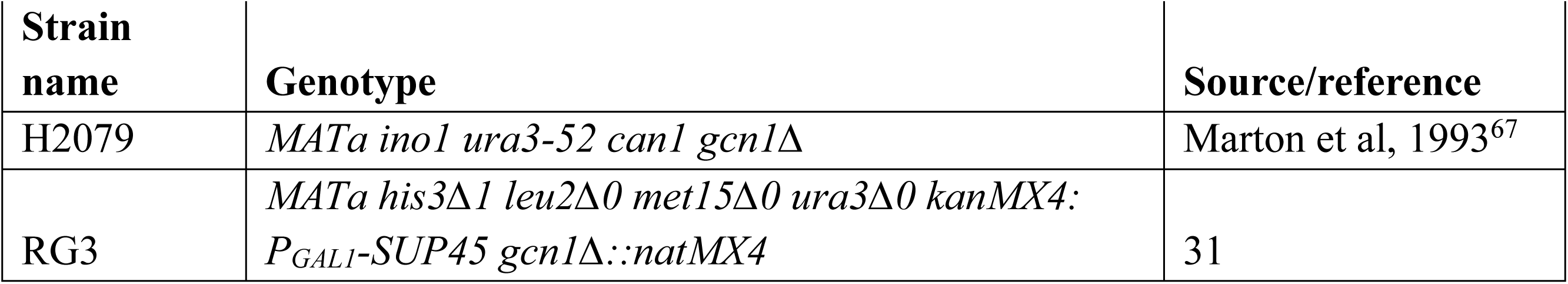
Yeast strains used in this study.

**Table S4.** Plasmids used in this study.

| Plasmid name | Description | Source/reference |
| --- | --- | --- |
| pRS316 | *URA3, CEN6* | Sikorski et al, 1989<sup>68</sup> |
| p2367 | *GCN1-myc* in pRS316 | Marton et al, 1997<sup>32</sup> |
| pRG20 | *gcn1-Δ746-805-myc* in pRS316 | This study |
| pRG21 | *gcn1-Δ1160-1176-myc* in pRS316 | This study |
| pRG22 | *gcn1-Δ746-805, Δ1160-1176-myc* in pRS316 | This study |

**Figure S1.**
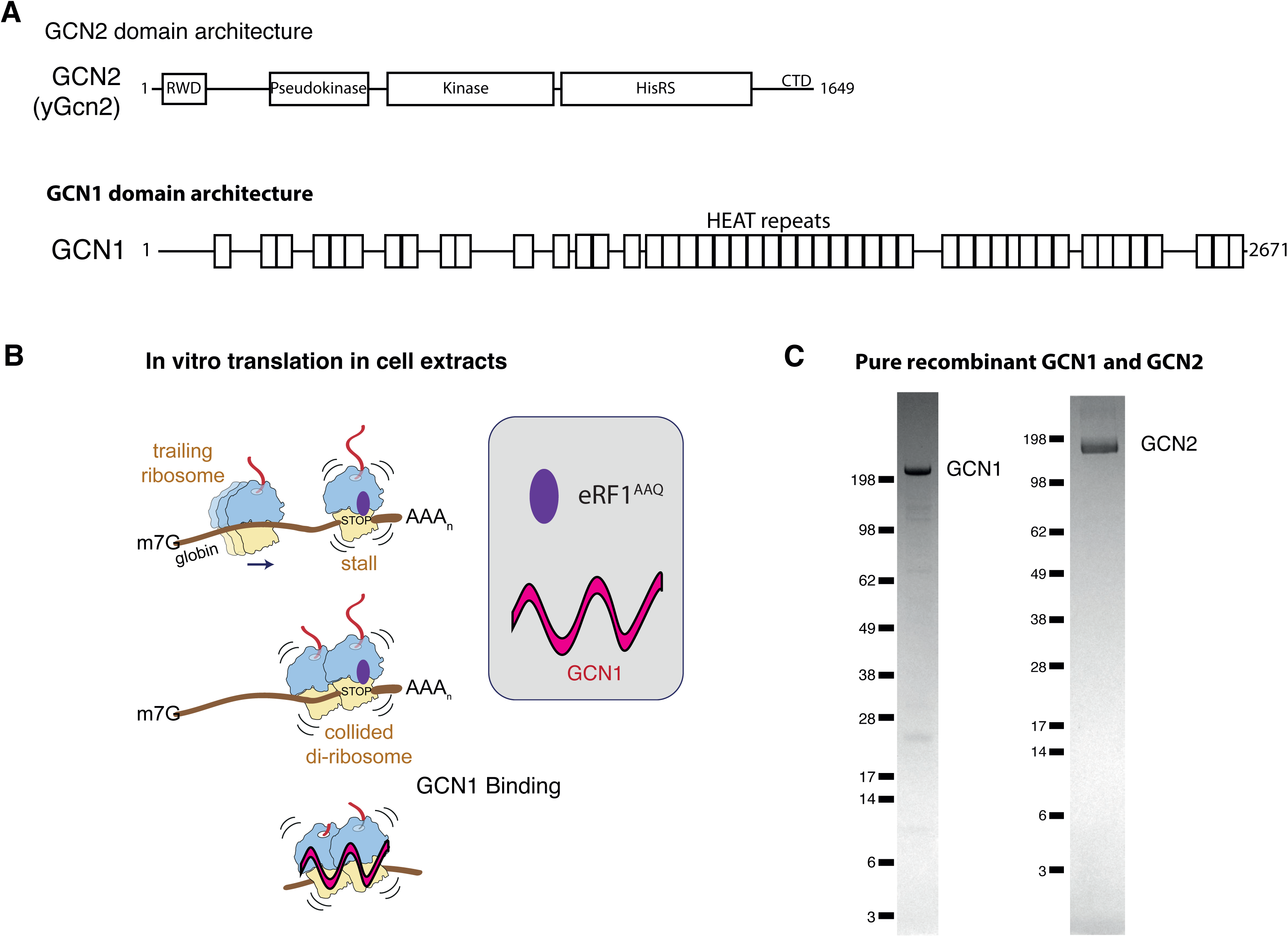
GCN protein-collided di-ribosome complex assembly. **(A)** Domain architecture of GCN2 and GCN1. **(B)** Schema depicting the strategy employed for assembling GCN1-collided di-ribosome complexes in vitro. Polysomes translating globin mRNA were stalled using dominant-negative mutant release factor eRF1^AAQ^ in the presence of GCN1, GCN2 and EDF1. **(C)** SDS-PAGE analysis of purified recombinant GCN1 and GCN2 expressed in Sf9 insect cells stained with InstantBlue Coomassie protein stain.

**Figure S2.**
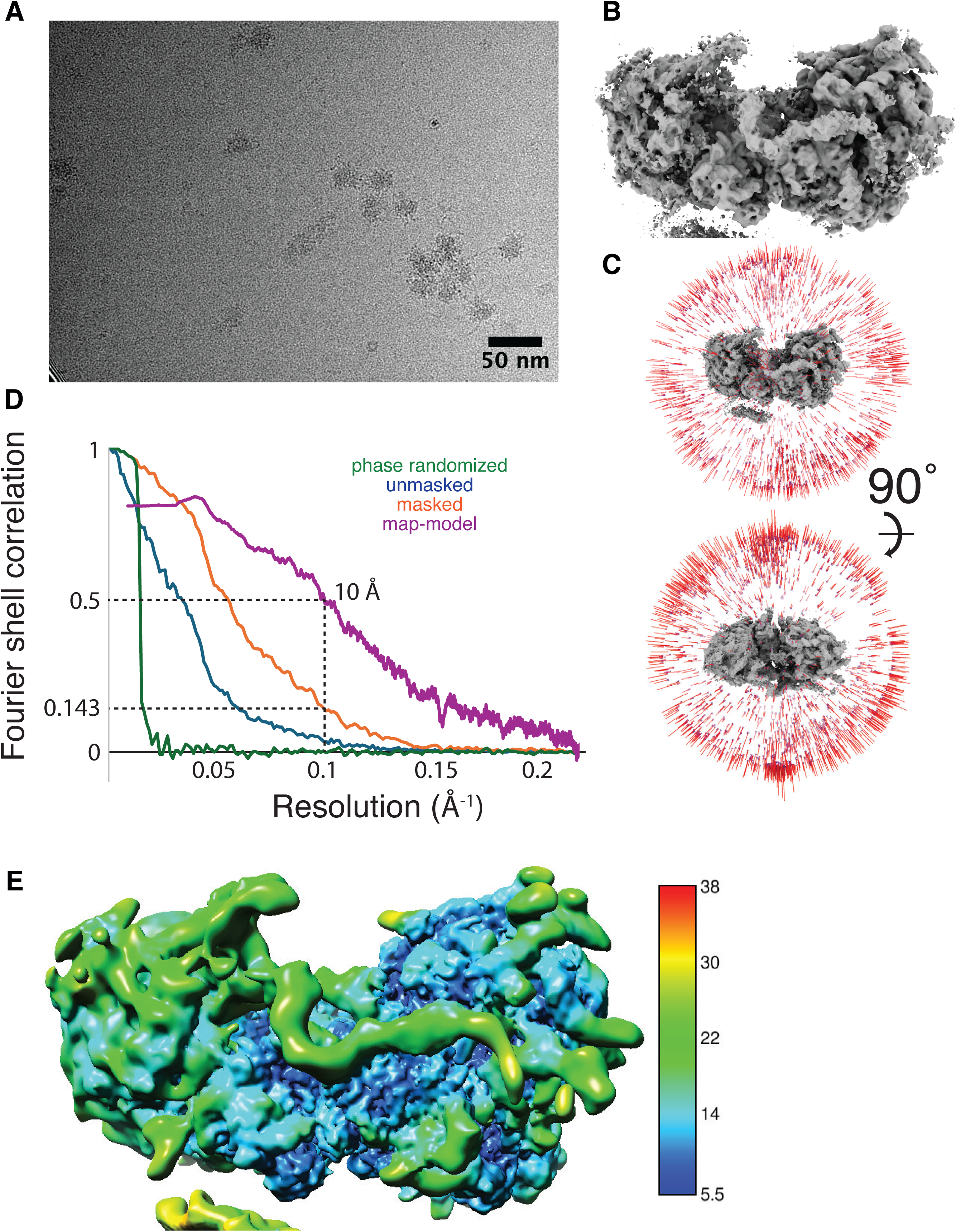
Structure determination of the GCN1-collided di-ribosome complex. **(A)** Representative micrograph, **(B)** final reconstructed map, **(C)** orientation distribution, **(D)** Fourier shell correlation (FSC) plots for the two independently refined, unmasked, dose-weighted half-maps, phase-randomized dose-weighted half maps, the final independently refined, masked, dose-weighted half-maps and the map-model FSC after model refinement are shown, **(E)** map filtered and colored according to local resolution.

**Figure S3.**
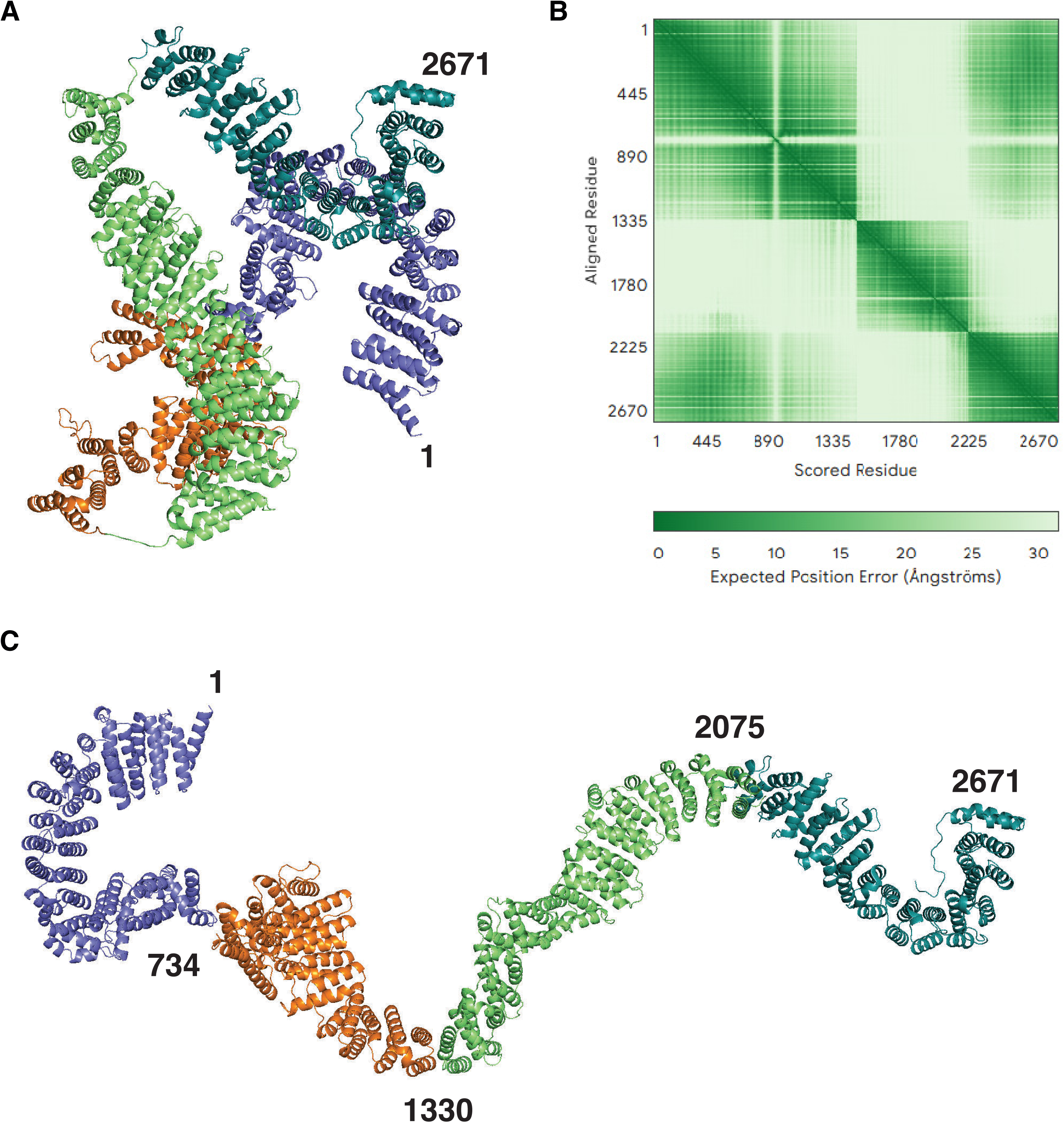
Alphafold3 prediction of GCN1 in the absence of the collided di-ribosome reveals a compact molecule. **(A-B)** Alphafold 3 prediction (A) and plot of expected position error in Å of the predicted structure (B). **(C)** Model of GCN1 bound to the collided di-ribosome used as scaffold in Phenix PredictandBuild. The domains were broken up based on the high confidence region boundaries along the diagonal in B and rigid-body docked into the cryo-EM density in Coot.

**Figure S4.**
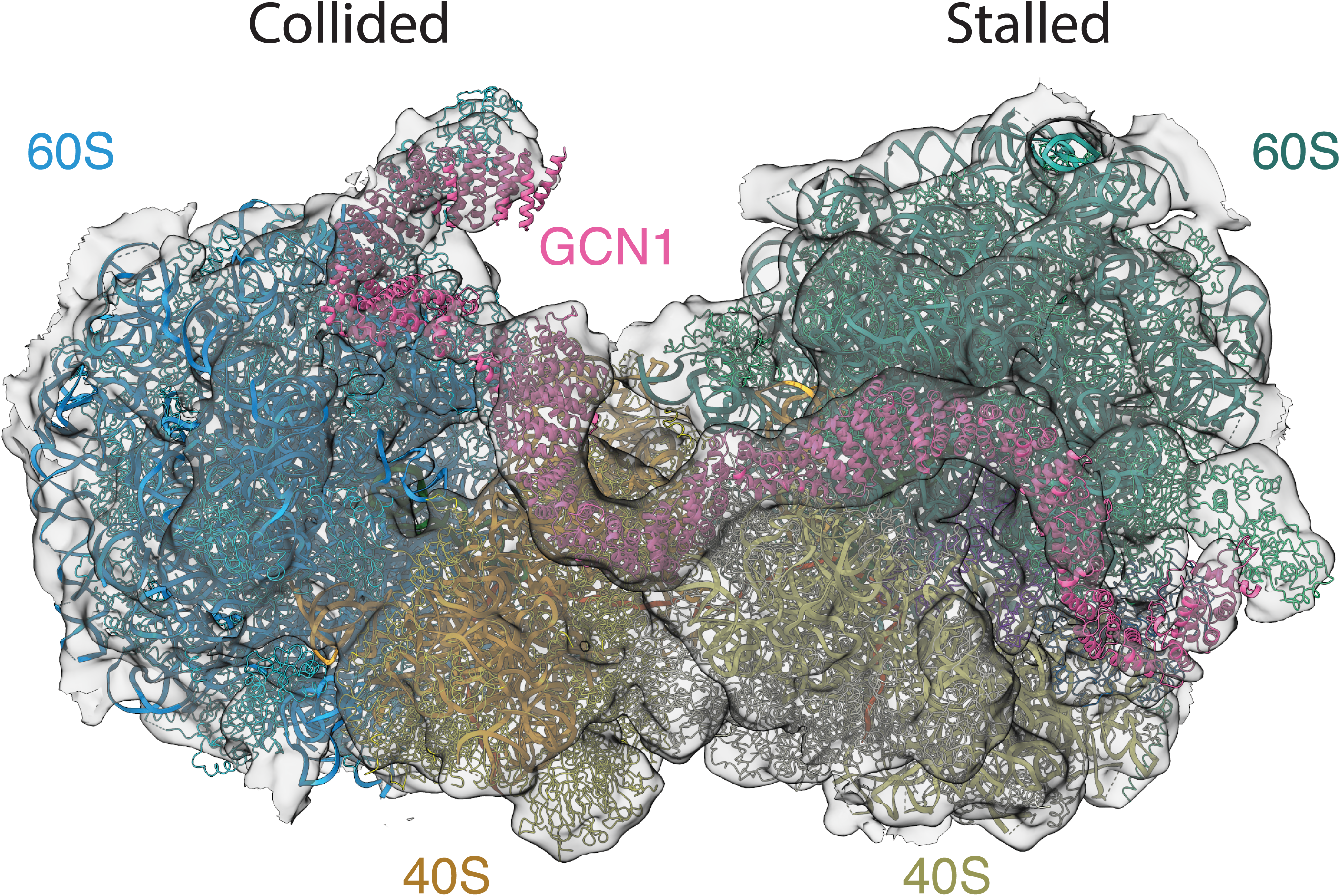
Map-to-model fit of the GCN1-collided di-ribosome structure. The 60S subunits are colored blue (collided) and teal (stalled). The 40S subunit is are colored orange (collided) and yellow (stalled). GCN1 is colored pink and the cryo-EM density is shown as a translucent gray surface. The figure was generated in UCSF Chimera 1.19.

**Figure S5.**
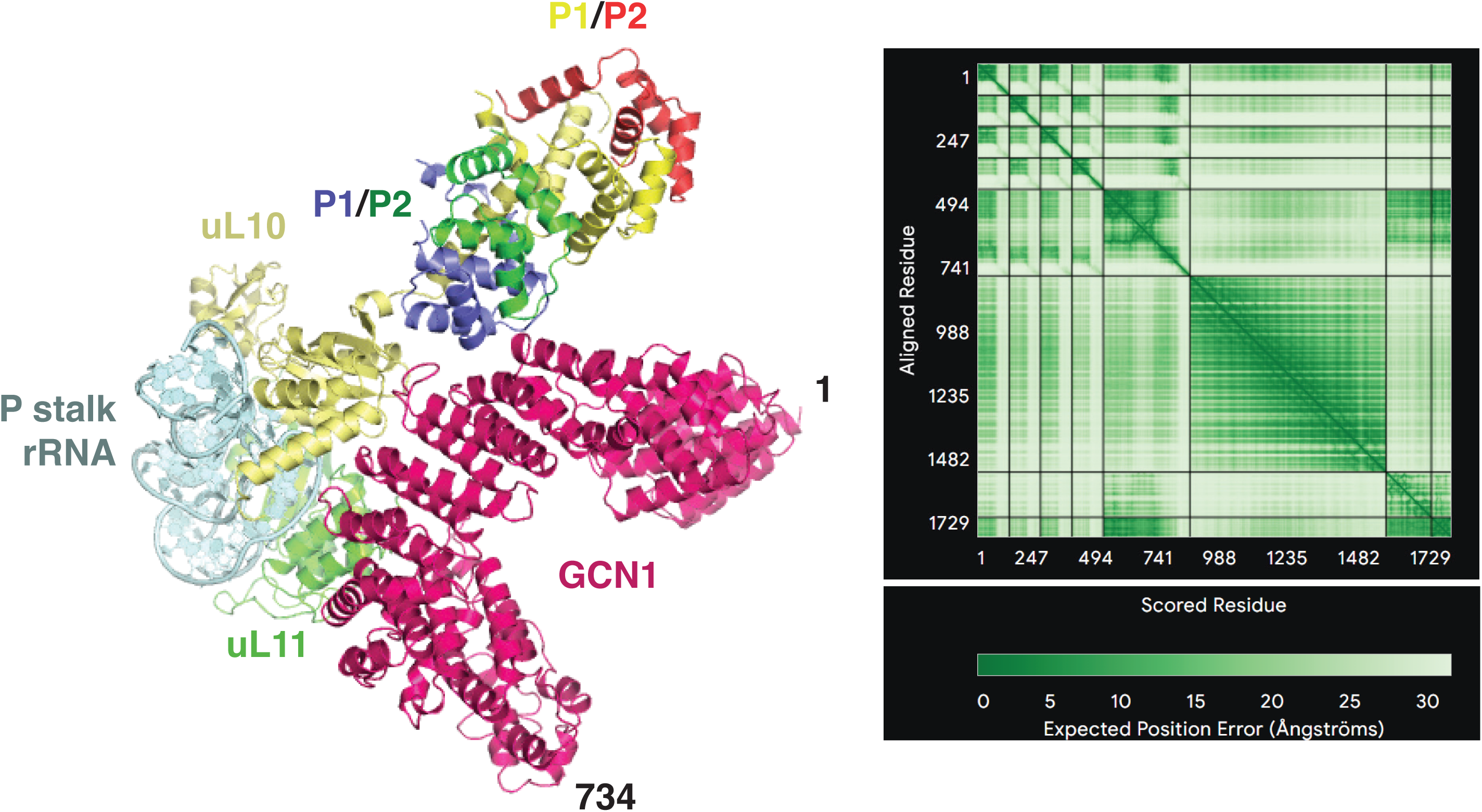
Alphafold3 prediction of the GCN1 N terminus interacting with the P stalk of the ribosome. Only residues 1-734 of GCN1 and nucleotides of the ribosomal 28S rRNA corresponding to the P stalk rRNA (residues 1484-1554 from *Oryctolagus cuniculus*) were included in the prediction. Expected position error in the predicted structure (in Ångströms) is shown on the right.

**Figure S6.**
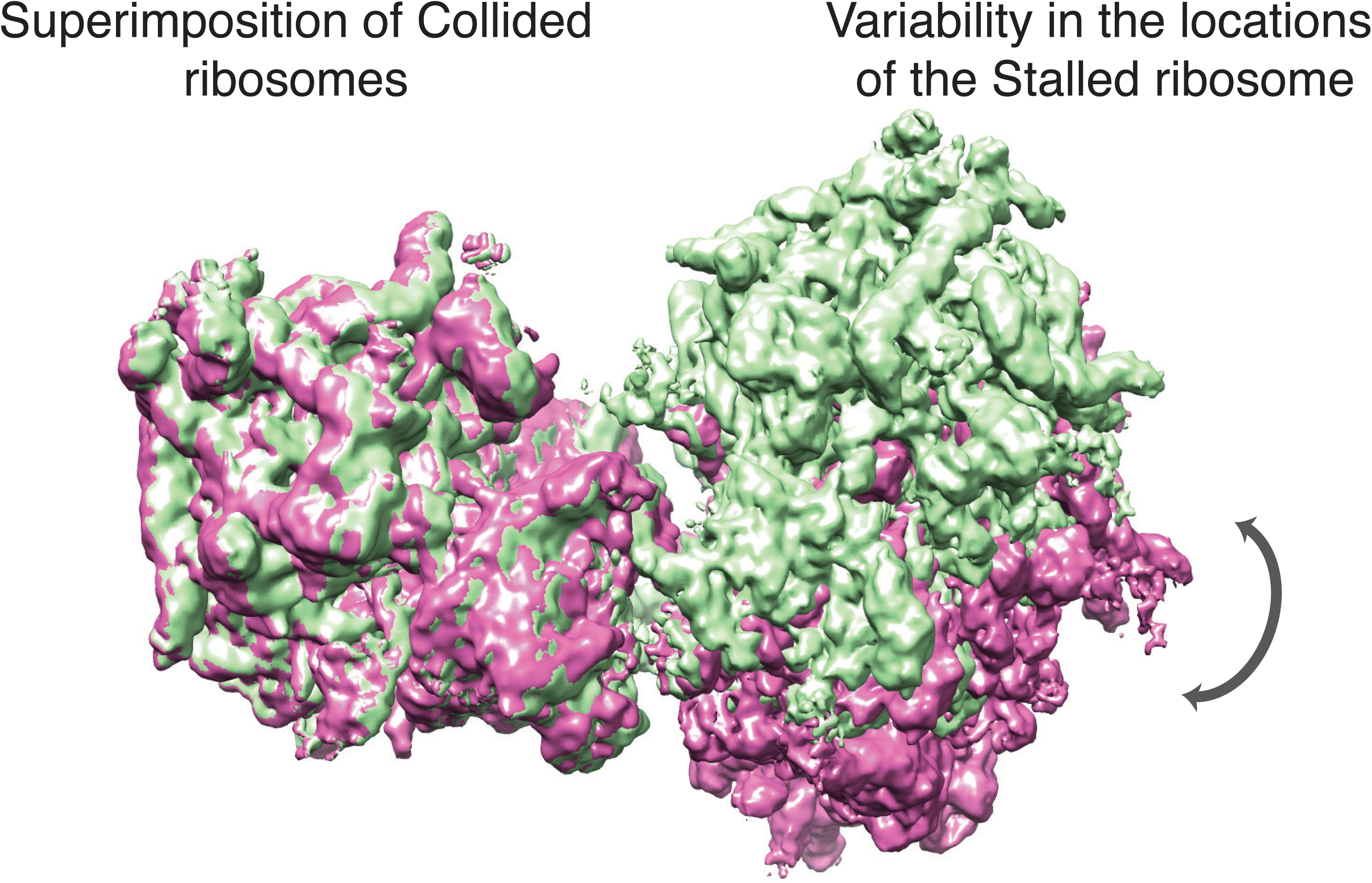
Flexibility of the collided di-ribosome when GCN1 is not bound. The flexibility of the stalled ribosome relative to a static collided ribosome was quantified by multi-body refinement and principal component analysis in RELION; particles were binned based on eigenvalues. Reconstructions of the top and bottom 10% of particles (corresponding to the maximum and minimum eigenvalues) are colored green and pink, respectively. These maps are superimposed on the collided ribosome to highlight conformational variability in the stalled ribosome between these particle subsets (indicated by the double arrow).

**Figure S7.**
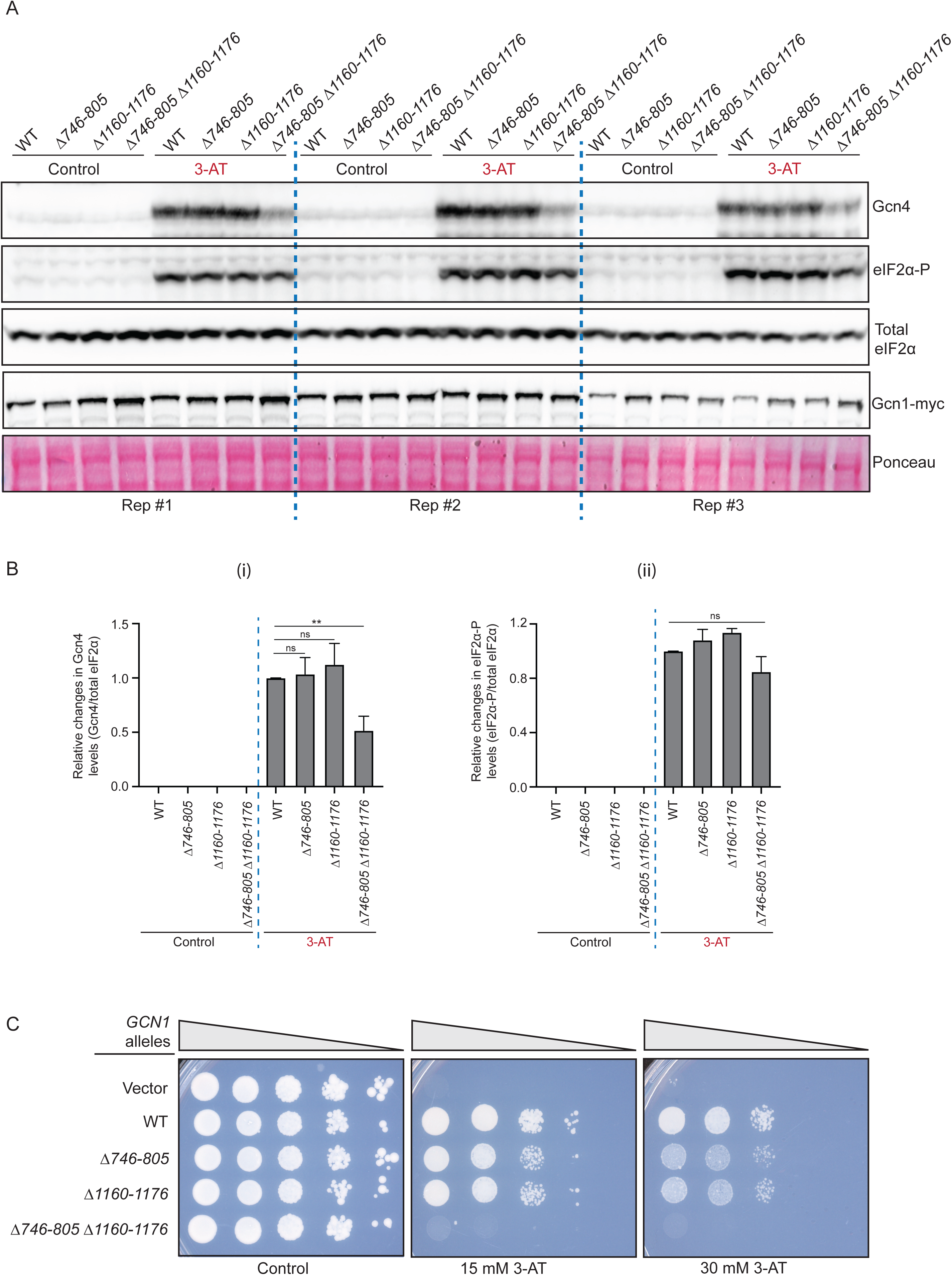
Gcn1 interaction with the 40S beak of the colliding ribosome is required for WT induction of Gcn4 in histidine-starved cells. **(A)** Immunoblot analysis of *gcn1*Δ strain (H2079) transformed with the indicated *GCN1-myc* alleles grown in SC-Ura containing 2% glucose (Control) or in SC-Ura-His containing 2% glucose and treated with 10 mM 3-AT for 1h (3-AT). The blot was probed with antibodies against the indicated proteins, with total eIF2α and Ponceau S staining shown as loading controls. **(B)** Immunoblot signals in (A) for Gcn4 and eIF2α-P were normalized to those for total eIF2α for all three biological replicates and used to calculate the means ± SD shown in the histograms for Gcn4 (i) and eIF2α-P (ii), with asterisks denoting statistical significance determined by a Student’s t-test: **p<0.01, ns-non-significant difference. **(C)** The same strains from (A) were grown in SC-Ura containing 2% glucose, 10-fold serially diluted, and spotted on the same solid medium (control), or on SC-Ura-His containing 2% glucose supplemented with 15 mM or 30 mM 3-AT, and incubated for 3-4 days at 30°C.

**Figure S8.**
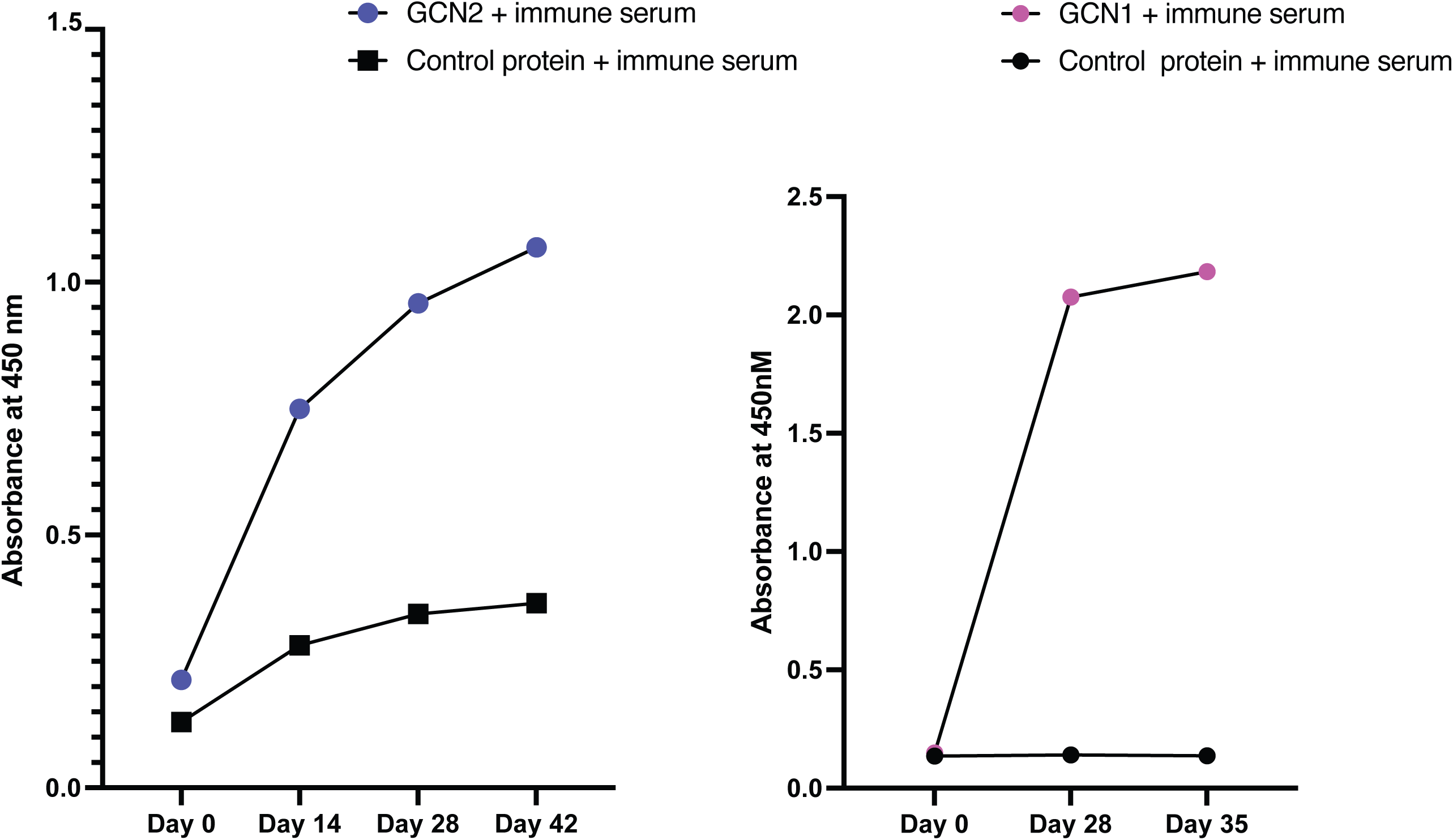
ELISA analysis of immune alpaca serum binding to biotinylated GCN1 and GCN2 antigens. Immune alpaca serum binding to biotinylated GCN1 and GCN2 was assessed by ELISA using biotinylated chicken lysozyme protein as an unrelated control protein. Absorbance of the TMB reaction product was measured at 450 nm.

**Figure S9.**
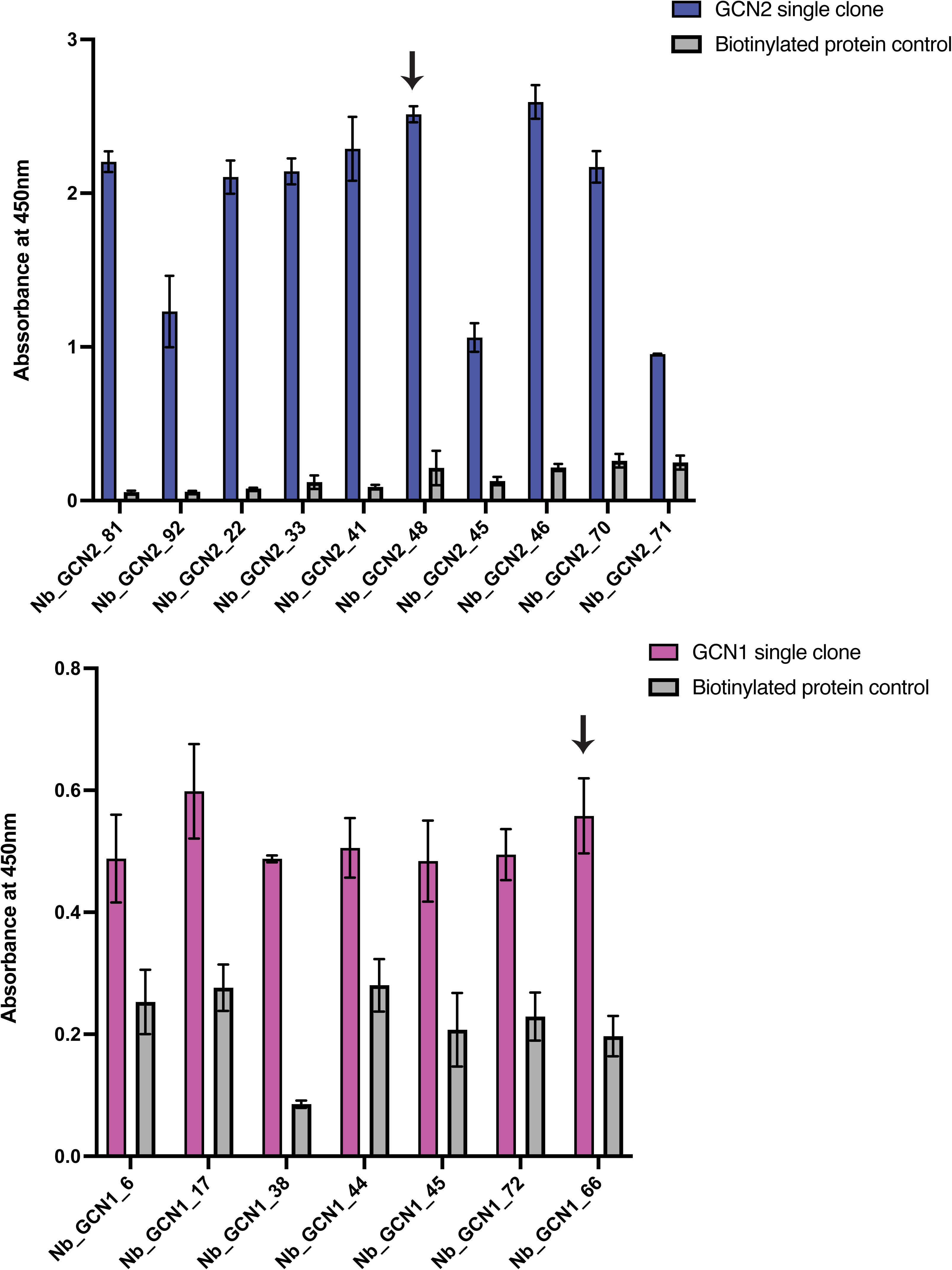
Monoclonal phage/periplasmic extract ELISA screening of individual nanobody clones against biotinylated GCN1 and GCN2 antigens. Individual nanobody clones were screened for binding to biotinylated GCN1 and GCN2 using biotinylated chicken lysozyme protein as an unrelated control protein. GCN2 Nb48 and GCN1 Nb66 (indicated by arrows) were selected for subsequent affinity measurements by fluorescence anisotropy and used in the proteomics experiment.

**Figure S10.**
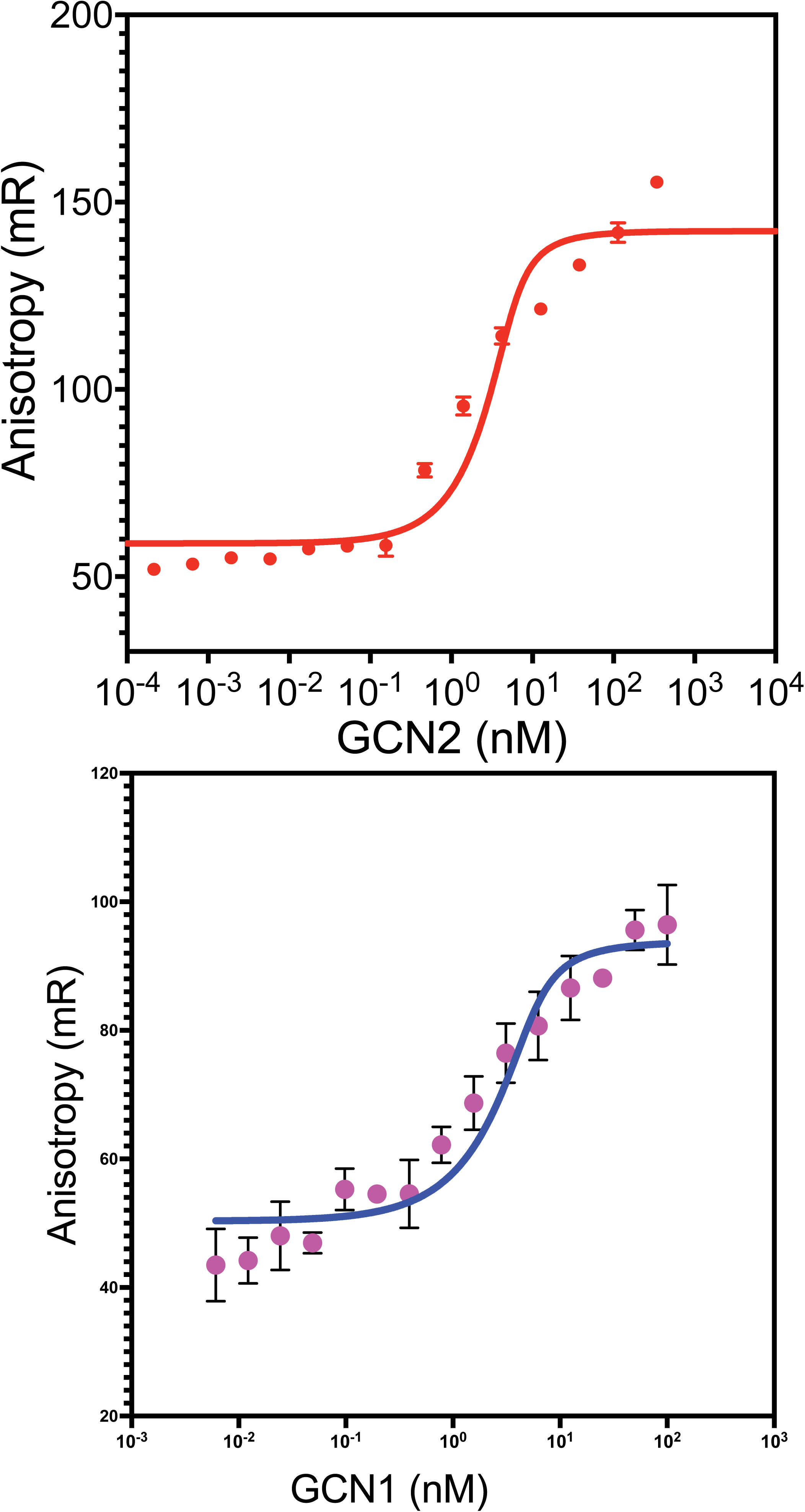
Affinities of GCN2/GCN1 nanobodies for GCN2/GCN1. Fluorescence anisotropy measurements with 5 nM Alexa Fluor 488-labelled GCN2 Nb48 or GCN1 Nb66 nanobodies titrating GCN2 or GCN1, respectively, after initial mixing or following a 20 min incubation. Apparent dissociation constants of 0.66 nM were obtained after 20 min for both GCN1 and GCN2, indicating high-affinity sub-nanomolar binding.

**Figure S11.**
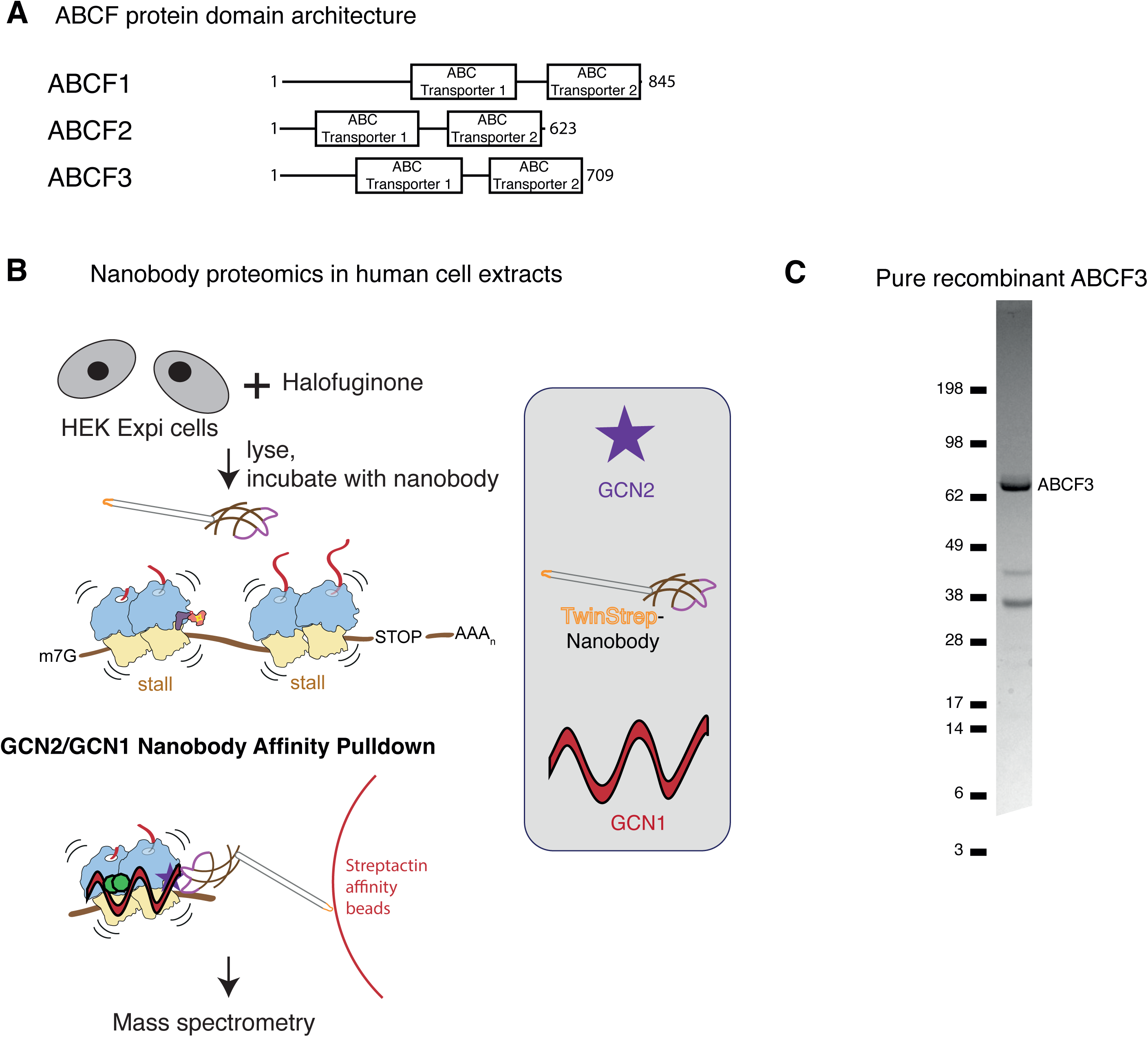
Identification of ABCF3 via affinity pulldown-mass spectrometry using α-GCN1 and α-GCN2 nanobodies. **(A)** Domain architecture of human ABCF proteins. **(B)** Schema depicting the strategy for identifying proteins that co-purify with GCN1 and GCN2. HEK Expi cells were treated with the prolyl-tRNA synthetase inhibitor halofuginone, lysed and incubated with recombinant Twin-Strep-tagged nanobody against either GCN2 or GCN1. Lysates were then incubated with Strep-Tactin XT sepharose resin and eluted with D-biotin. **(C)** SDS-PAGE analysis of purified recombinant ABCF3 expressed in Sf9 insect cells and stained with InstantBlue Coomassie protein stain.

**Figure S12.**
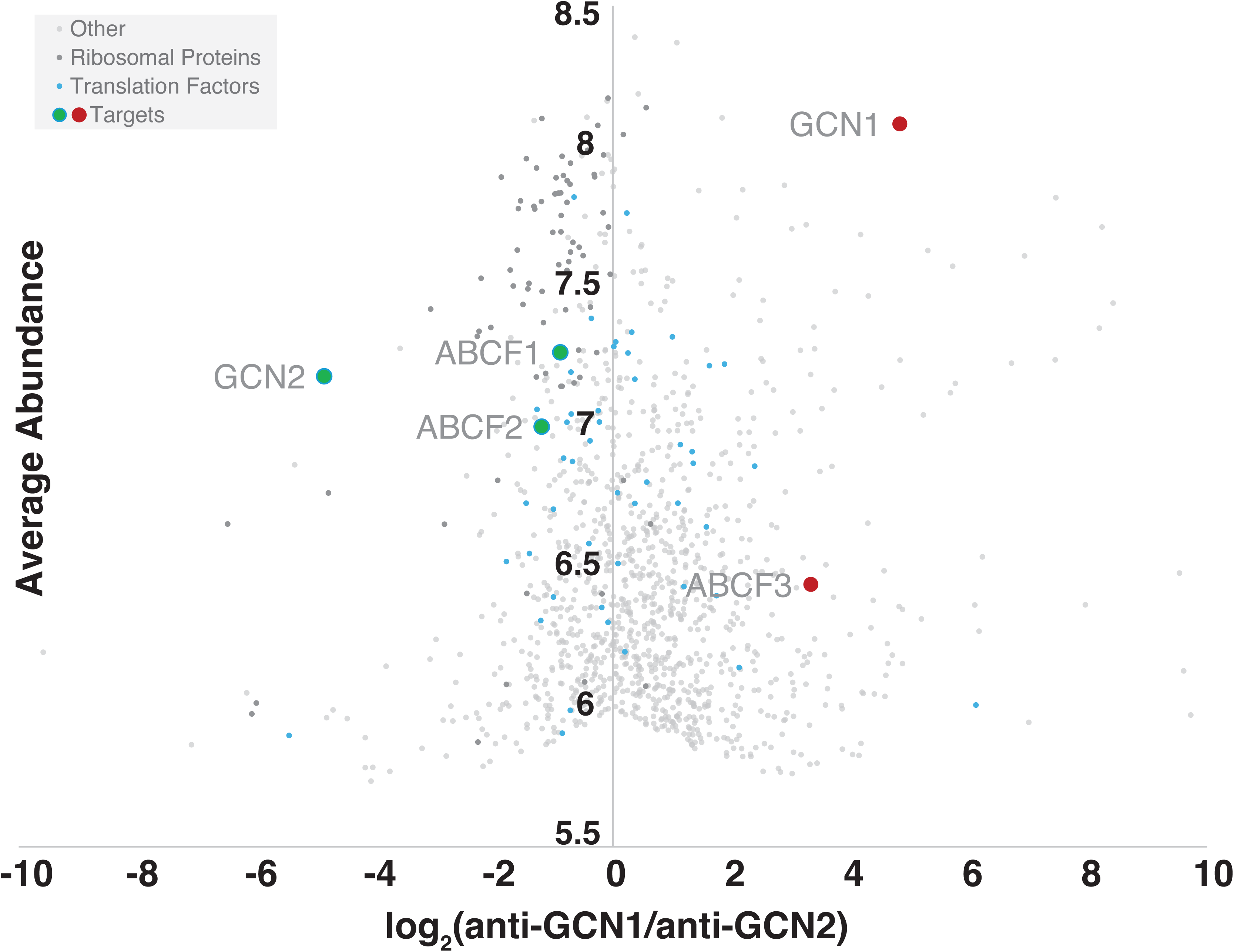
Identification of the human ortholog of yeast GCN20. Lysates from Expi293 cells treated with halofuginone for 30 minutes were incubated with immobilized anti-GCN1 or anti-GCN2 nanobody and the eluates were subject to mass spectrometry. Bland-Altman plot comparing hits from the anti-GCN1 nanobody with hits from the anti-GCN2 nanobody. X-axis - log_2_ (anti-GCN1/anti-GCN2), Y-axis - average abundance. Note that the Y-axis is truncated from 5.5 - 8.5 for clarity and proteins not seen in both samples are eliminated by limiting the X-axis to [-10, 10]. Ribosomal proteins are in dark grey, translation factors are colored light blue and proteins known to be involved in the amino acid starvation response are in green (anti-GCN2 nanobody) or red (anti-GCN1 nanobody). All other proteins are shown in light grey. All three human paralogs ABCF1-3 were pulled down by both nanobodies but ABCF3 was more effectively enriched by the anti-GCN1 nanobody while ABCF1 and 2 were enriched better by the anti-GCN2 nanobody.

## Notes

### Competing Interest Statement

V.R. is a founder and shareholder of RNAvate Private Ltd., an RNA therapeutics company. The goal of RNAvate is to deliver circular RNAs into cells for therapeutic expression of specific genes. The work here has no direct bearing on the interests of RNAvate. Moreover, this work is of purely academic interest and no patent has been filed as a result of this work. All other authors declare no competing interests.

